# Computation-guided redesign of promoter specificity of a bacterial RNA polymerase

**DOI:** 10.1101/2022.11.29.518332

**Authors:** Xiangyang Liu, Anthony T. Meger, Thomas Gillis, Jonah O’Mara Schwartz, Balendra Sah, Robert Landick, Srivatsan Raman

## Abstract

The ability to regulate genetic circuits and metabolic pathways is central to cellular control. The existing toolkit is predominantly comprised of local transcription regulators that are unsuitable for exerting control at a global genome-wide scale. Bacterial sigma factors are ideal global regulators as together they direct the RNA polymerase to thousands of transcription sites. Here, we redesigned the promoter specificity of the *E. coli* housekeeping sigma factor, sigma-70, toward five orthogonal promoter targets not recognized by the native sigma-70. These orthogonal sigma-70 factors were developed by screening a pooled library of computationally designed variants of the -35 DNA recognition helix, each tailored to a specific target promoter. In the redesigned sigma factors new target-specific interactions facilitate new promoter recognition. Activity of the top performing redesigned sigma-70s varied across the promoter targets and ranged from 17% to 77% of native sigma-70 on its canonical active promoter. These orthogonal sigma factors represent a new suite of regulators for global transcriptional control.

## INTRODUCTION

Synthetic biology aims to control cellular behavior through synthetic pathways and circuits. These pathways and circuits depend on orthogonal gene regulation to selectively insulate them from host processes while sharing central dogma resources and machinery with the host.(1–5) In bacteria, native gene regulation is governed at the local and global levels.(6–8) At the local level, individual genes and operons are controlled by transcription factors that are activated by specific metabolites or environmental cues. These local regulators (i.e., small-molecule inducible transcription factors) have been co-opted and engineered to build a large toolkit of orthogonal gene regulatory components for synthetic circuits and pathways.(9–12) At a global level, the RNA polymerase (RNAP) is directed to hundreds to thousands of loci simultaneously by a battery of sigma factors to maintain homeostasis and respond to environmental changes.(13) In contrast to local regulators, efforts to engineer RNAP to exert control at a global level remain poorly developed. Although some genes or operons can be regulated from synthetic promoters with non-native regulation, large swathes of the bacterial genome, especially genes involved in core metabolic processes, require native regulation as they are often layered in complex networks that cannot be easily disentangled.(3,14) Orthogonal regulation in these genes can be introduced by modifying RNAP promoter specificities without disrupting native regulation.(15) With emerging technologies to synthesize genomes from scratch, genome-scale orthogonal gene regulation could be programmed into cells.(16,17) Such a system could have different RNAP promoter specificities encoded in different regions of the genome to compartmentalize them. Thus, redesigning the RNAP is a powerful but underexplored route to engineer orthogonal genetic regulation.

The sigma subunit of the bacterial RNAP governs the interaction of the RNAP with the promoter.(18–20) The bacterial RNAP is a multi-subunit molecular machine composed of alpha, beta, omega, and sigma subunits. The alpha, beta, and omega subunits form the core RNAP enzyme responsible for RNA synthesis using DNA as a template and ribonucleotides as the substrate.(21,22) To initiate transcription, the core RNAP binds to a sigma factor which directs the RNAP to specific promoters.(19,20,23,24) Bacteria express several types of sigma factors that can recognize and bind specific promoters in response to cellular signals and environmental conditions.(24–26) Of the seven types of sigma factors in *E. coli*, sigma-70 is the dominant as it is responsible for expressing housekeeping genes, and the other sigma factors are only transiently expressed during stress, require additional activator binding sites, and/or are expressed at much lower concentrations. (27–32) Prior strategies to engineer RNAP have used heterologous sigma factors from other hosts or constructed chimeras of native and non-native sigma factors to engineer new RNAP promoter specificities.(33–35) However, heterologous sigma factors are often toxic to the host, and chimeric sigma factors are not orthogonal, limiting the extent to which they may be used in synthetic systems.(36,37) Additionally, the use of heterologous and chimeric sigma factors is strain-dependent.(33) Despite the attractiveness of the sigma factor as a target for orthogonal genetic regulation, we do not have well-developed strategies to engineer them.

In this study, we redesigned the *E. coli* sigma-70 to recognize and transcribe from five orthogonal promoters that are transcriptionally inactive to wild-type sigma-70, using computation-guided design and pooled cell-based screens. We used Rosetta to create a customized library of computationally redesigned variants of the domain of *E. coli* sigma-70 that interacts with each promoter target. Fluorescence-activated cell sorting followed by deep sequencing of the sigma libraries revealed the sequence determinants of promoter specificity and how they changed for different targets. We found that recognition of new promoters occurs through a combination of highly conserved residues that likely make polar interactions with DNA backbone and target specific adaptations. We identified orthogonal sigma variants for each of the five targets whose activities ranged from 17-77% of wildtype sigma-70 on a highly active canonical *E. coli* promoter. Our workflow is generalizable and can be applied to alternative sigma factors in *E. coli* and sigma factors from other host organisms. The redesigned sigma factor-promoter pairs constitute an important tool for engineering orthogonal genetic regulation at the genome level.

## MATERIALS AND METHODS

### Rosetta design

All protein-DNA modeling calculations were performed with the Rosetta v3.9 macromolecular modeling program as described previously.(38) The crystal structure of *E. coli* sigma-70 in complex with the canonical -35 element (PDB: 4YLN) was used as the scaffold for redesign. The -35 element was substituted to each of the five target promoter elements: TTCATC, GGAACC, CCGCCG, GCTACC, and CCCCTC. We performed a combinatorial mutagenesis scan of sigma-70 positions R584, E585, R586, R588 and Q589 to generate all single, double, triple, and quadruple mutants. The stability of each resulting protein-DNA interface was calculated by taking the average score across 10 Rosetta optimized structures. The 1000 sigma-70 variants with the highest affinity (i.e. lowest protein-DNA interface score) on each -35 target were selected for experimental testing. For targets TTCATC, GGAACC, and CCGCCG, we generated an additional test set of 1000 variants with binding energies nearest to native sigma-70 and the canonical -35 element (-26.0 REU).

### Library preparation and cloning

A promoter library was ordered via Integrated DNA Technology and amplified by hybridizing with a reverse primer (hybridization protocol: 95°C for 3min, 55°C for 1min, 72°C for 1min). The product was subsequently cleaned up using PCR cleanup kit (Omega Biotek) and stored in dH_2_O. Backbone preparation started with PCR amplification of plasmid pXL-9. For Rosetta designed sigma variants, 110-base pair (bp) single strand DNA oligo pool containing Rosetta designed sigma fragments were ordered from Agilent. Oligo design features unique priming regions at 5’ and 3’ for individual libraries complemented with BsaI recognition sites and cutting sites to enable Goldengate cloning. 10ng of the oligo pool was used in the PCR reaction to amplify individual libraries (PCR protocol: 95°C for 3min, 98°C for 20sec, 55°C for 15sec, 72°C for 8sec, cycling from step 2 for 20 cycles, 72°C for 30sec). PCR product was subsequently cleaned up using PCR cleanup kit (Omega Biotek) and stored in dH_2_O. Backbone preparation started with PCR amplification of plasmid SC101_LacI_WTsigma containing a WT sigma-70 driven by pLacO promoter. Amplified backbone bared the corresponding BsaI binding and cutting sites, enabling scarless cloning. Backbone PCR was cleaned up and subject to sequential digestion of DPN1 and BsaI_HF_V2 (New England Biolab) to remove the template DNA and expose sticky overhangs. The digested backbone was further incubated with Antarctic Phosphatase (New England Biolab) to remove 5’ and 3’ phosphates. The removal of those phosphates prevents backbones from circularizing and creating false positive transformants. The Goldengate reaction was done as described on New England Biolab. Briefly, 300ng of the backbone was combined with 70ng of the library in a 20µL reaction. The mix was incubated at 37°C for 1hr and 65°C for 5min. Cooled reaction mix was dialyzed on 0.025 µm filter in dH_2_O for 1hr at room temperature. The dialyzed sample was collected and stored at -20°C.

### Transformation of promoter and redesigned sigma-70 libraries

Library transformation was conducted using electrocompetent DH10β *E. coli* (New England Biolab). 2µL of the assembled library reaction mix was transformed with 25µL cells and recovered with 1mL SOC medium at 37°C for 1hr. Multiple logs of dilutions were plated on appropriate antibiotic plate to measure transformation efficiency. 4mL of LB and antibiotics were added to ensure proper growth selection. Cultures were grown overnight before being stored in 25% glycerol at -80°C. For non-library transformations, 10ng DNA was added directly to 25µL of cells and plated on agar plates containing antibiotics to confer selection. Co-transformations were conducted by first growing cells containing one plasmid in LB medium with appropriate antibiotics overnight. 1:50 back dilution was then done into 3mL LB without antibiotics. Cells were allowed to grow until reaching an OD_600nm_ of 0.6 before being chilled on ice. Chilled cells were spun down at 5000xg and thoroughly washed with ice cold dH_2_O twice. The final cell pellet was suspended in 25µL of ice cold dH_2_O. Electroporation was then repeated similarly to regular plasmid transformation. Transformed cells were plated onto agar plates with appropriate antibiotics overnight at 37°C before storage at 4°C.

### Induction and fluorescence measurement

Multiple colonies were picked from agar plates and inoculated into 150µL LB medium in a 96-well plate containing appropriate antibiotics. Cells were allowed to grow shaking (900rpm, multi-well plate shaker, Southwest Scientific) at 37°C for around 3hr or until OD_600nm_ reached 0.6. Cultures were back inoculated into two technical replicates of fresh LB medium with appropriate antibiotics at 1:20 dilution factor. After around 3hr or upon reaching an OD_600nm_ of 0.3, IPTG (at final concentration of 5µM) was added to one of the two technical replicates. Cells were incubated for 5hr before fluorescence measurement. Fluorescence measurements were conducted using a multi-well platereader (HTX Biotek). Excitation and emissions wavelengths of 485nm and 528nm (± 20) were used for all measurements. Measurements were normalized by dividing fluorescence intensities by the OD_600nm_ before subtracting blank well fluorescence (cells carrying no GFP). Fold-improvement was calculated by dividing normalized the IPTG induced fluorescence of sigma-70 variants by the IPTG induced fluorescence of WT sigma-70 containing cells using the same target promoter sequence. Average fold improvement was calculated from at least 3 biological replicates.

### Fluorescence activated cell sorting

Around 25µl of reporter transformed libraries carrying a target promoter upstream of GFP were inoculated into 3mL LB kan/spec-50 (kanamycin and spectinomycin at 50µg/mL) from glycerol stock and grown overnight at 37°C in a shaking incubator. Cultures were inoculated into two separate 150µL fresh LB kan/spec-50 in a 96-well plate at 1:50 dilution. Fresh cultures grew shaking at 37°C for 3hr or until reaching an OD_600nm_ of 0.3. IPTG was added to a final concentration of 5μM to one technical replicate for induction. After the addition of IPTG, cultures were grown for 5hr before being chilled on ice. Chilled cultures were added to ice cold phosphate buffer saline (PBS) at 1:50 ratio and mixed well before being put back onto the ice. Cell sorting was conducted using a Sony SH800 cell sorter (Sony Biotech) (condition: FCS threshold=2500, 50% PMT on GFP, ultra-purity, round 8k events/second). Three independent replicates were sorted. Briefly, cells in PBS were flowed through first to capture the distribution. Under 5µM IPTG induced condition, top 10% cells were sorted out (at least 50k events). Sorted populations were flowed through the sorter to capture sorting efficiency (percentage of cells fall back into the sorted gate). Sorted cells with less than 80% efficiencies were sorted using the same gate again to improve the purity. Sorted cells were recovered in 1mL LB medium shaking at 37°C. 100µL (1:10 dilution) of cells were plated onto LB agar plates with appropriate antibiotics. The rest were growing overnight in 3ml LB with appropriate antibiotics before being miniprepped into plasmid. Promoter library sorting was done by simply isolating 5% of cells with lowest fluorescence.

### Variant identification using next generation sequencing (NGS)

A pair of primers upstream and downstream of the mutated sigma-70 region were used to add partial Illumina adaptors. Priming sites were at least 20 base pair away from the variable regions to ensure good quality base calling during NGS. Amplicon size was 120 base pair. Amplicons were sequenced on Amplicon-EZ Miseq service (Genewiz) with at least 50k reads for each sigma library. Sequenced libraries were processed with PEAR pair-ended merger (quality threshold set to 35). Merged reads were processed with custom script to identify variants. Briefly, sigma-70 variable regions were extracted and converted into amino acids with corresponding frequency. Reads with frequencies of less than 5 were removed. Variant frequency was normalized against total reads to get percentages. To compute individual variant’s enrichment, a variant’s percentage under 10µM IPTG induction was divided by that of the condition without IPTG. Sequence logos for variants with highest enrichment were generated using ISA-tools.(39)

### Clonal identification

To identify functional sigma-70 variants, 100 colonies were sampled from the sorted populations. Highly-functional (defined by its fold improvement level) colonies were measured using the above protocol for plate reader measurements, and high performing colonies were subsequently identified using Sanger sequencing. Unique high-performing variants were then inoculated in 3 mL kan/spec LB media and grown overnight. Following overnight growth, plasmids from these variants were extracted using the protocol for plasmid mini-prep. Following plasmid extraction, plasmids from these variants were re-transformed into DH10β *E. coli* in order to normalize for variations in the *E. coli* genome of sorted cells that may have led to false positives in GFP transcription during clonal testing. Using three biological replicates, the re-transformed sigma-70 variants were measured once again using the induction protocol from initial clonal screens. High-performing sigma-70 variants were measured alongside cells containing WT sigma-70 and the orthogonal promoter sequences to ensure that fold-improvement scores are consistent across measurements.

## 5′ Rapid Amplification of cDNA Ends (5′ RACE)

E. coli cells harboring either wild-type or engineered promoters and sigma 70 variants were grown in 5 mL of LB broth containing the appropriate antibiotics. Cultures were induced at OD600 ∼0.3 by adding 1 mM IPTG and grown for ∼6 hours to allow GFP expression. Total RNA was extracted from 3mL of culture using TRIzol reagent (Invitrogen) followed by isopropanol precipitation following manufacturer’s directions.

For 5′ Rapid Amplification of cDNA Ends (5′ RACE), the FirstChoice RLM-RACE kit (Ambion) was used with minor modifications. Three micrograms of RNA were treated with Tobacco Acid Pyrophosphatase (TAP) at 37°C for 1 hour before ligating the 5′ RACE adapter. cDNA synthesis was then performed according to the manufacturer’s protocol. The cDNA was amplified using adapter and GFP specific primers using one Taq Polymerase (NEB) (PCR protocol: 94°C for 30 sec, 94°C for 20sec, 55°C for 20sec, 68°C for 30sec, cycling from step 2 for 30 cycles, 68°C for 5min). The PCR product was purified and submitted for sanger sequencing (Functional Biosciences). The resulting read was aligned to the original sequence and the TSS was determined by where the original sequence ends, and the adapter sequence begins.

## RESULTS

### The partitioned functional domains of sigma-70 enable targeted redesign of promoter specificity

As the “housekeeping” regulator of transcription in *E. coli*, sigma-70 performs three functions. First, sigma-70 must recognize the consensus -10 (TATAAT) and -35 (TTGACA) promoter elements. The -10 element is recognized by conserved domains 2 and 3, and the -35 element is recognized by the helix-turn-helix motif of domain 4 (domain 1 is unstructured) (**Fig. 1a**).(40,41) Second, sigma-70 is responsible for recruiting the core RNA polymerase complex to the promoter. The interface between RNA polymerase and sigma-70 is extensive (>8000 Å^2^) as it spans the entire length of domains 2-4.(42) The third function of sigma-70 is to mediate the melting of DNA near the -10 region, which is a critical step for initiating transcription. To operate as a global regulator of cellular transcription, sigma-70 must perform each of these functions with high efficiency and fidelity.

**Fig. 1.**
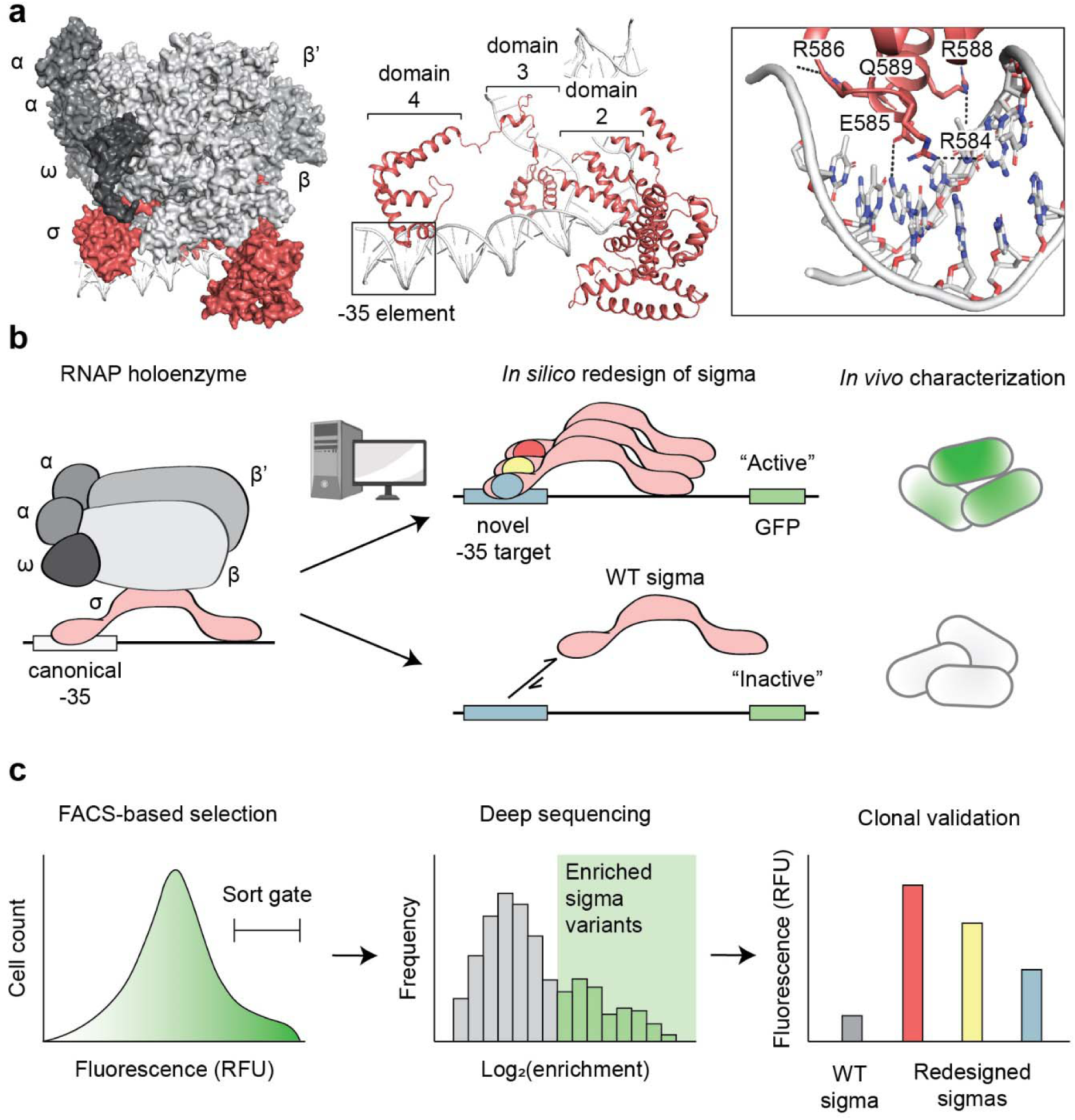
Engineering sigma-70 variants with novel promoter specificity. (**a**), Structure of the *E. coli* transcription initiation complex (PDB ID: 4YLN). sigma-70 (red) bound to the -10 and -35 DNA elements (gray, cartoon representation) and RNAP core complex (gray, surface representation), composed of α (two copies), β, β^’^, and ω subunits (left). Canonical -35 element recognized by the helix-turn-helix motif of sigma-70 domain 4 and -10 element recognized by domains 2 and 3 (middle). Orientation and h-bond contacts of residues R584, E585, R586, R588, and Q589 of the domain 4 recognition helix in the -35 major groove (right). (**b**), Workflow for altering sigma-70 promoter specificity *in silico* and characterizing function *in vivo*. Cartoon schematic of the RNAP holoenzyme with WT sigma-70 bound to the canonical - 35 element (left). Replacement of the canonical -35 with a novel target and computational redesign of the domain 4 recognition helix to engineer sigma-70 variants with orthogonal promoter specificities (middle). *In vivo* functional characterization using a GFP gene placed downstream of the promoter variants containing novel -35 targets (right). (**c**), Platform for high-throughput selection and characterization of redesigned sigma-70 variants. FACS to isolate GFP positive (i.e., functional variants) (left), deep sequencing to identify enriched variants (middle), and clonal testing of redesigned sigma-70s to validate function (right).

To engineer orthogonal sigma variants, we must alter promoter specificity while preserving interactions with the core RNA polymerase complex and those required for DNA melting. Therefore, we aimed to restrict the design to key residues of the helix-turn-helix motif of domain 4 to alter promoter specificity through the -35 DNA element. These include residues R584, E585, R586, R588, and Q589 (**Fig. 1a, box**). In the structure of the *E. coli* transcription initiation complex (PDB ID: 4YLN), the positively charged R586 and R588 form polar interactions with the negatively charged phosphate backbone to likely promote general DNA affinity.(41) Specific interactions with the nucleotide bases are formed between R584:Gua-31 and E585:Ade-34 in the major groove of the -35 element. Although oriented towards the major groove, Q589 does not directly interact with the -35 element. Instead, Q589 is sterically packed between residues E585, R586 and R588 and, likely plays a role in properly orienting these critical residues within the recognition helix. Given their structural roles, we hypothesized that redesigning these five key positions could engineer orthogonal promoter specificity of sigma-70.

Our design workflow begins with identifying -35 element mutants that are not recognized by wildtype sigma-70. These mutants became recognition targets for the computational redesign of the major groove helix residues of wildtype sigma-70 (**Fig. 1b**). The top 1000 ranked designs were synthesized as chip-based oligonucleotide library and evaluated in pooled fashion using a cell-based GFP screen. In this assay, high GFP fluorescence signifies a specificity switch of sigma-70, as the variant must recognize the orthogonal -35 element to initiate transcription of GFP. To identify specificity switches from the pooled screen, we compared the distribution of variants between sorted and presorted states by deep sequencing (**Fig. 1c**). Clonal screens were then used to validate several successful redesigns for each targeted -35 element.

### Synthetic promoter modifications enhance sigma-70 dependence on the -35 element

As mentioned above, sigma-70 utilizes a two target DNA recognition system, with domain 4 recognizing the canonical -35 element, and domains 2 and 3 recognizing the canonical -10 element. Due to this two-target dependence, engineering an orthogonal sigma factor would require the redesign of multiple sigma-70 DNA binding domains and their respective consensus DNA sequences. However, by artificially increasing dependence of DNA recognition on the -35 DNA element, redesign is simplified to a one domain-one target problem. There are several advantages to engineering orthogonality through the -35 element rather than the -10 element: (1) -35 recognition is mediated by a single domain of sigma-70, (2) interactions between the helix-turn-helix motif of domain 4 with the -35 major groove are structurally well understood, and (3) interactions between sigma-70 and the -10 element are much more complex as transcription initiation is mediated by DNA melting near this region. Thus, we sought to increase sigma-70’s dependence on the -35 element, and then engineer sigma-70 variants that recognize orthogonal -35 targets.

We first placed a gene encoding GFP downstream of a strong constitutive *E. coli* promoter with canonical -35 ‘TTGACA’ and -10 ‘TATAAT’ sequences (apFab71)(43) on a “reporter plasmid” to use fluorescence as a measure of transcriptional activity. As expected, a negative control lacking both canonical -10 and -35 promoter elements resulted in very low fluorescence relative to the apFab71 promoter (**Fig. 2b**). We generated a randomized (NNNNNN) library of the -35 sequence to assess the dependence on the -10 site. Moderate fluorescence was observed in the population of cells expressing the randomized -35 library, suggesting that the canonical -10 element alone is partially sufficient to recruit sigma-70. To attenuate the -10 dependence and increase the -35 dependence, we aimed to identify a promoter variant with high transcriptional activity with the canonical -35 sequence, but significantly lower activity when paired with a disrupted -10 sequence. To this end, we characterized a small subset of nucleotide deletion variants (Δ(-11), Δ(-16,-7), Δ(-8), Δ(-7)) near the -10 region of apFab71. The Δ denotes deletion of a nucleotide at a specific site within the -10 site. We quantified the -35 dependence of each -10 deletion variant by comparing the fluorescence ratio of the deletion variant with a clonal canonical -35 and the randomized -35 population. As desired, all four apFab71 promoter variants retained -35 dependence (i.e., lower fluorescence in the randomized condition) and displayed approximately a 10-fold reduction in fluorescence relative to the native apFab71 promoter with a randomized -35 sequence (**Fig. 2a**). The Δ(-7) promoter exhibited the greatest -35 dependence with a 330-fold difference in fluorescence between the canonical and randomized -35 sequences. Thus, the Δ(-7) promoter was selected as the starting construct for redesigning the promoter specificity of sigma-70.

**Fig. 2.**
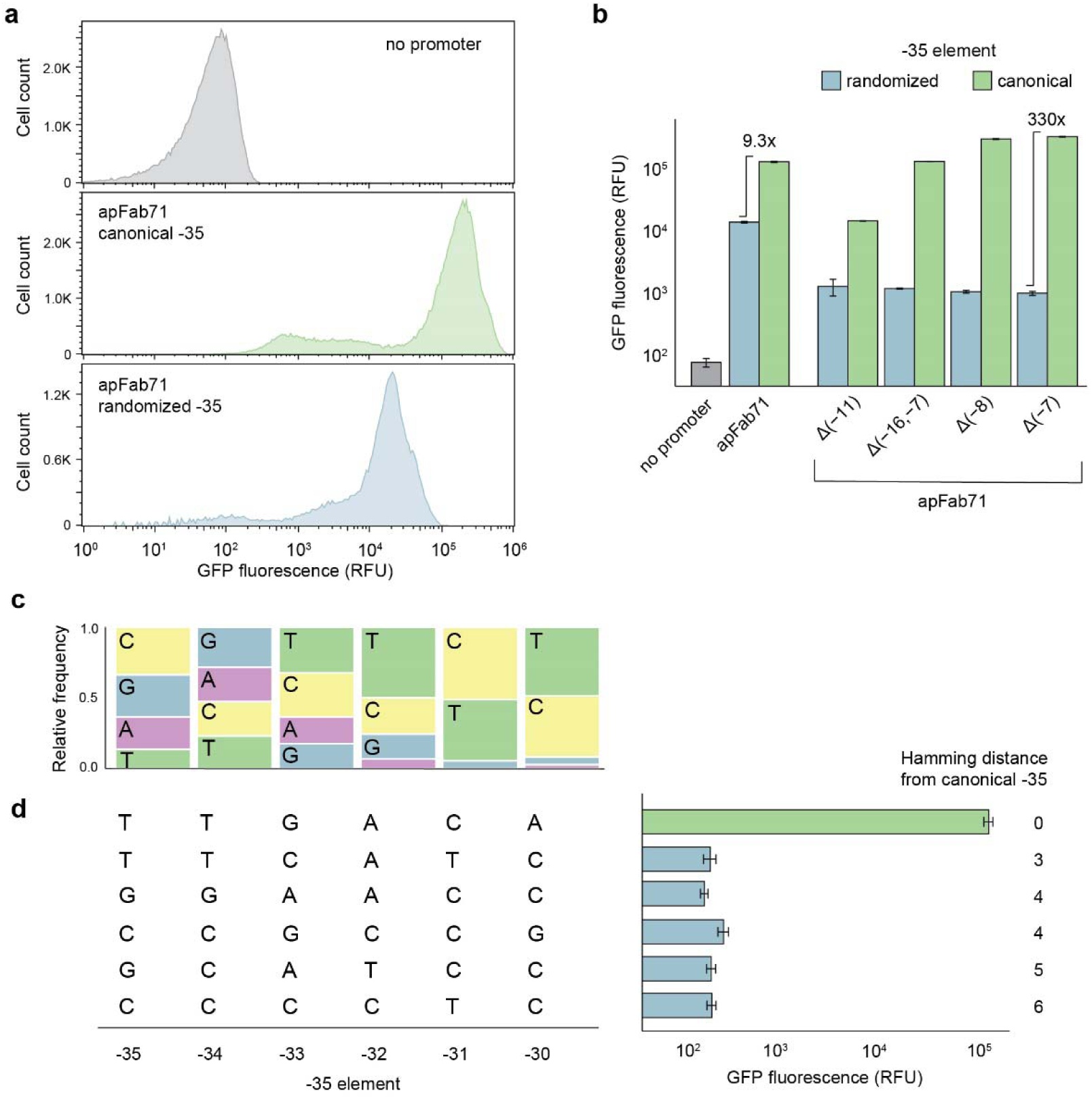
Sigma-70 dependence on the -35 element and selection of orthogonal promoter targets. (**a**), Fluorescence distributions of *E. coli* cells expressing native sigma-70 and containing plasmid constructs with a GFP gene placed downstream of no promoter (gray), the strong constitutive apFab71 promoter (green), or the apFab71 promoter with a randomized -35 element (blue). (**b**), Median flow cytometry measured fluorescence of promoter variants containing nucleotide deletions (Δ) near the -10 region, and either a canonical or randomized -35 element. Fold-change between canonical and randomized -35 states was used to evaluate -35 transcriptional dependence. (**c**), Sequence logos (-35 region) of 74 unique clonal isolates with low GFP fluorescence (<10^3^ RFU) (**Supplementary Table 1**). Clones were screened in 96-well format from the apFab71 Δ(-7) randomized -35 library. (**d**), Selected -35 DNA targets for engineering orthogonal sigma-70 variants. Nucleotide sequences (left) and median fluorescence (right) using native sigma-70. Error bars denote the standard deviation of replicate flow cytometry experiments (n≥2). RFU, relative fluorescence units. OD, optical density measured at 600nm.

### Endogenous *E. coli* sigma-70 displays very weak activity on selected -35 targets

We next sought to identify a set of five promoter targets orthogonal to the canonical -35 element recognized by endogenous *E. coli* sigma-70. From the library of randomized -35 sequences of the apFab71 Δ(-7) promoter, we used a high-throughput 96-well screening approach to isolate and characterize promoter variants not recognized by native sigma-70. We isolated 74 unique -35 variants with a 500-fold weaker sigma-70 transcriptional activity (GFP fluorescence <10^3^ RFU) relative to apFab71 (**Supplementary Table 1**). Comparing the sequences of these low activity -35 variants did not reveal a strong pattern suggesting that there are many unique ways to disrupt sigma-70 promoter recognition (**Fig. 2c**). Weak conservation was observed at Cyt-31 of the canonical -35 (TTGACA), but other positions deviated from the canonical sequence which is also reflected by their median Hamming distance of 5 (**Supplementary Table 1**). From these low activity -35 variants, we selected five diverse target sequences for engineering sigma-70 variants with novel specificities: TTCATC, GGAACC, CCGCCG, GCTACC, and CCCCTC. These promoter targets all displayed low transcriptional activity (500- to 900- fold lower fluorescence than apFab71), comparable to the no promoter control, and vary in their composition and Hamming distances from the canonical -35 sequence (**Fig. 2d**). Due to their diversity, we hypothesized that these five -35 sequences would present unique design challenges and demonstrate the breadth of our redesign capabilities.

### Rosetta calculations reveal the sigma-70 sequence preferences for each promoter target

To engineer sigma-70 variants that recognize each of the five -35 targets (TTCATC, GGAACC, CCGCCG, GCTACC, and CCCCTC), we first performed large-scale combinatorial mutagenesis of residues R584, E585, R586, R588, and Q589 in the helix-turn-helix (HTH) motif of domain 4 using the Rosetta macromolecular modeling suite.(44–46) We excluded I587 because it is located along the interface of the HTH and makes no DNA contacts. The design workflow is as follows. We replace the canonical -35 sequence (TTGACA) of the sigma-70:DNA co-crystal structure (PDB: 4YLN, **Fig. 1a**) with a target -35 sequence that we experimentally validated was unable to initiate transcription using native sigma-70 (**Supplementary Table 1**). Then, we perform an *in silico* scan to evaluate the stability of the protein-DNA complex for all possible single, double, triple, and quadruple variants (total ∼724,000 sigma-70 variants) of the sigma-70 recognition helix (Fig. 1a, boxed, middle panel). Calculations for all ∼724,000 sigma-70 variants against the five target -35 elements were performed on a high-throughput computing (HTC) system, requiring approximately 400,000 computing hours (2-3 weeks real-time). Computed protein:DNA interface scores for the ∼724,000 sigma variants in complex with the promoter targets were used to curate designs for experimental testing (**Fig. 3a, Supplementary Fig. 1a**). Lower interface scores indicate a more stable complex. Interestingly, differences between median interface scores among sigma-70 variants revealed a hierarchy of promoter preference of the sigma-70 scaffold (TTGACA < TTCATC < GGAACC < CCGCCG < GCATCC < CCCCTC) with interface scores decreasing with increasing Hamming distance from the consensus -35 sequence, TTGACA.

**Fig. 3.**
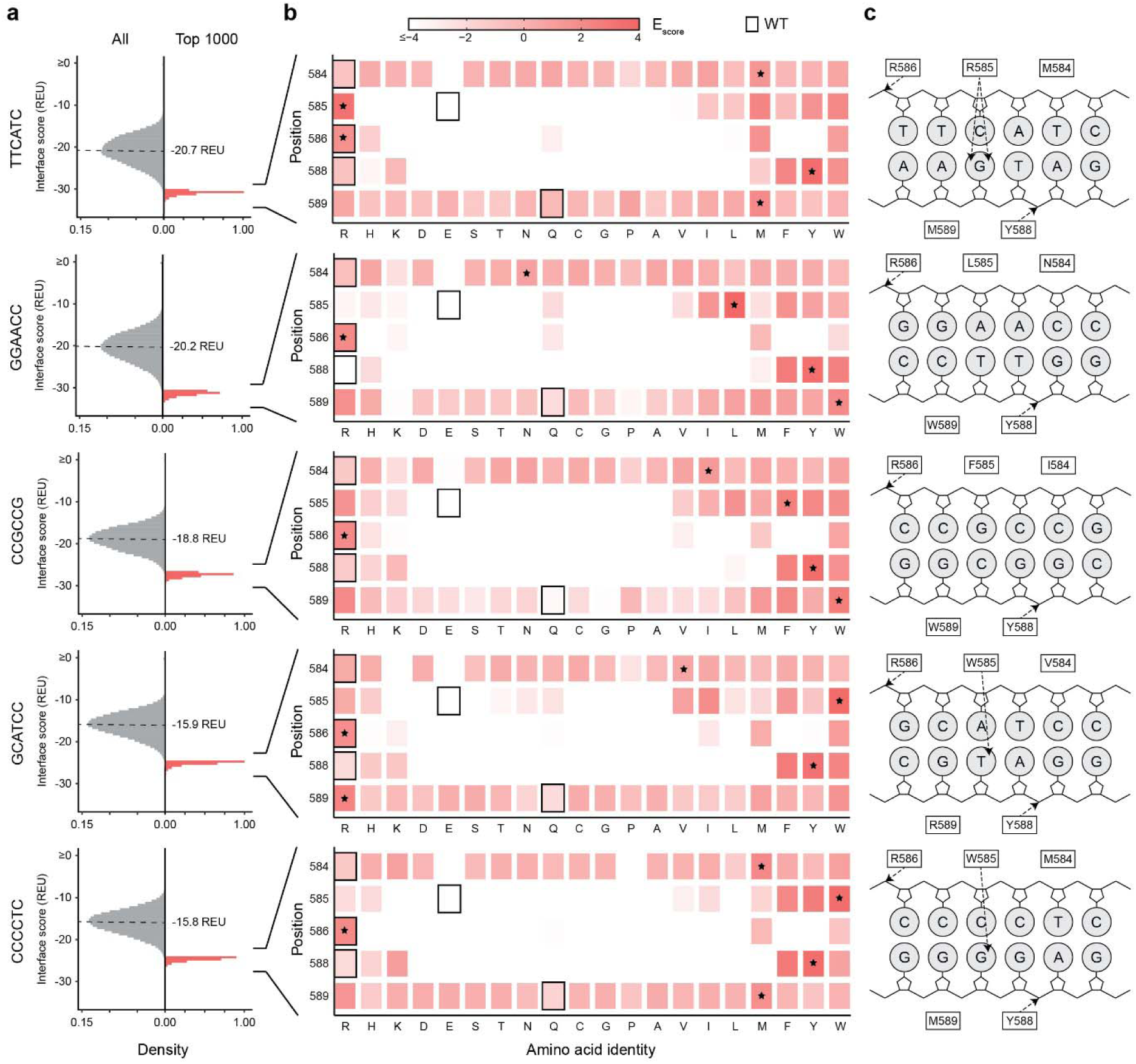
Rosetta guided design of sigma-70 variants for each promoter target. (**a**), Interface scores of all (gray) or 1000 top scoring (red) sigma-70 variants in complex with each -35 DNA target. Dashed lines indicate the median interface scores of all single, double, triple, and quadruple combinatorial variants of sigma-70 positions 584, 585, 586, 588, and 589 modeled with Rosetta. (**b**), Position-specific amino acid enrichment scores (red gradient) among selected top scoring sigma-70 variants. WT identity (boxed outline) and most enriched amino acid (*) at each mutable position. (**c**), Cartoon schematic showing H-bonds formed between each enriched sigma-70 consensus sequence and -35 DNA target in the Rosetta structural models.

This suggests a smooth binding landscape of sigma-70 DNA recognition where incremental mutations from the canonical -35 sequence decrease binding strength. To assess sigma-70 sequence preferences with each promoter, we computed position-specific amino acid enrichment scores (E_score_) for the top 1000 scoring interfaces (**Fig. 3b, Supplementary Fig. 1b**). We observed that substitutions at positions 584 and 589 did not have an impact on the protein-DNA interface scores suggesting that these are indirect contacts that likely do not drive affinity. Nonetheless, the most enriched residue at both positions was not the wildtype amino acid and was distinct for each promoter site. This suggests that though these may be indirect contacts, they could play a role in promoter specificity. In contrast, positions 585, 586 and 588 show subtle, but distinct preferences for different binding sites. Enrichment heat maps indicate sigma-70 positions 585, 586, and 588 undergo stringent selection across all -35 targets. These positions directly contact DNA in the crystal structure of native sigma-70 in complex with the canonical (TTGACA) -35 element (**Fig.1a**). As expected, positively charged and/or large hydrophobic amino acids are generally preferred at these positions, which can facilitate highly favorable hydrogen bonding, electrostatic and/or van der Waals interactions with the DNA. For all promoters, arginine and tyrosine were the most enriched amino acids at positions 586 and 588, respectively. Both residues make non-specific hydrogen bonds with the sugar-phosphate backbone of DNA (**Fig. 3c, Supplementary Fig. 1c**). Clear differences in amino acid preference were evident for position 585. For the -35 target TTCATC, arginine was the most enriched amino acid, which forms two hydrogen bonds with Gua-35 in the Rosetta structural models. Similarly, a single hydrogen bond is formed by tryptophan at position 585 with Gua-35 or Thy-35 for promoters CCCCTC and GCATCC, respectively. In contrast, bulky hydrophobic residues phenylalanine and leucine make van der Waals contacts with the CCGCCG and GGAACC promoters. Because positions 584 and 589 do not directly interact DNA, substitutions at these positions had less distinct effects on the Rosetta computed interface scores.

The redesigned sigma-70 variants were ranked by binding energy, and the highest affinity 1000 variants for each target -35 element were selected for experimental testing. In addition, for targets TTCATC, GGAACC, and CCGCCG, we selected 1000 variants with binding energies comparable to native sigma-70 and the canonical -35 element (-26.0 REU), as this affinity may be optimal for DNA recognition and promoter release during transcription initiation. For GCATCC and CCCCTC, the “highest affinity” and “WT-like” sets were identical because the binding energy distributions of these targets were less favorable (**Fig. 3a**). We synthesized these variant libraries on an IPTG-inducible plasmid expression system using oligonucleotide chip synthesis and one-pot cloning (see **Library preparation and cloning)**.

### Isolation and identification of sigma-70 variants with novel promoter specificities

To synthesize the computationally curated library of sigma-70 designs for each promoter target, we used chip-based oligonucleotides and one-pot cloning. The library of sigma-70 designs for each target was independently cloned into an expression plasmid and placed under IPTG control. Each library was transformed into *E. coli* harboring their cognate reporter plasmid i.e., GFP gene placed downstream of the respective target -35 element. Due to the “housekeeping” role of sigma-70, overexpression of a sigma-70 variant can competitively sequester the RNAP core enzyme to inhibit cell viability. This toxicity challenge has been reported in previous works using heterologous sigmas and domain swaps.(33–35) To identify an expression level with minimal toxicity, we first varied the IPTG inducer concentration in a clonal growth assay. Minimal growth deficiency was observed using 5μM IPTG, and thus, this concentration was used for all subsequent experiments (**Supplementary Fig. 2**).

To isolate and enrich high activity sigma-70 redesigns, we sorted the high fluorescence subpopulation of each library in the IPTG induced condition. Comparison of the fluorescence distributions of cell populations in the induced and uninduced states show distinctly higher fluorescence in the induced condition for all five target promoters (**Fig. 4a, Supplementary Figs. 3,4a**). This suggested that some sigma-70 redesigns may indeed be able to successfully transcribe from an orthogonal -35 sequence. The presorted and sorted libraries were deep sequenced to identify sigma-70 redesigns that were enriched after selection. Approximately 10-15% of all sigma-70 variants within each 1000-member library were enriched (**Fig. 4b, Supplementary Fig. 4b**). Taken together, these results validate our *in silico* redesign approach and high-throughput screening platform.

**Fig. 4.**
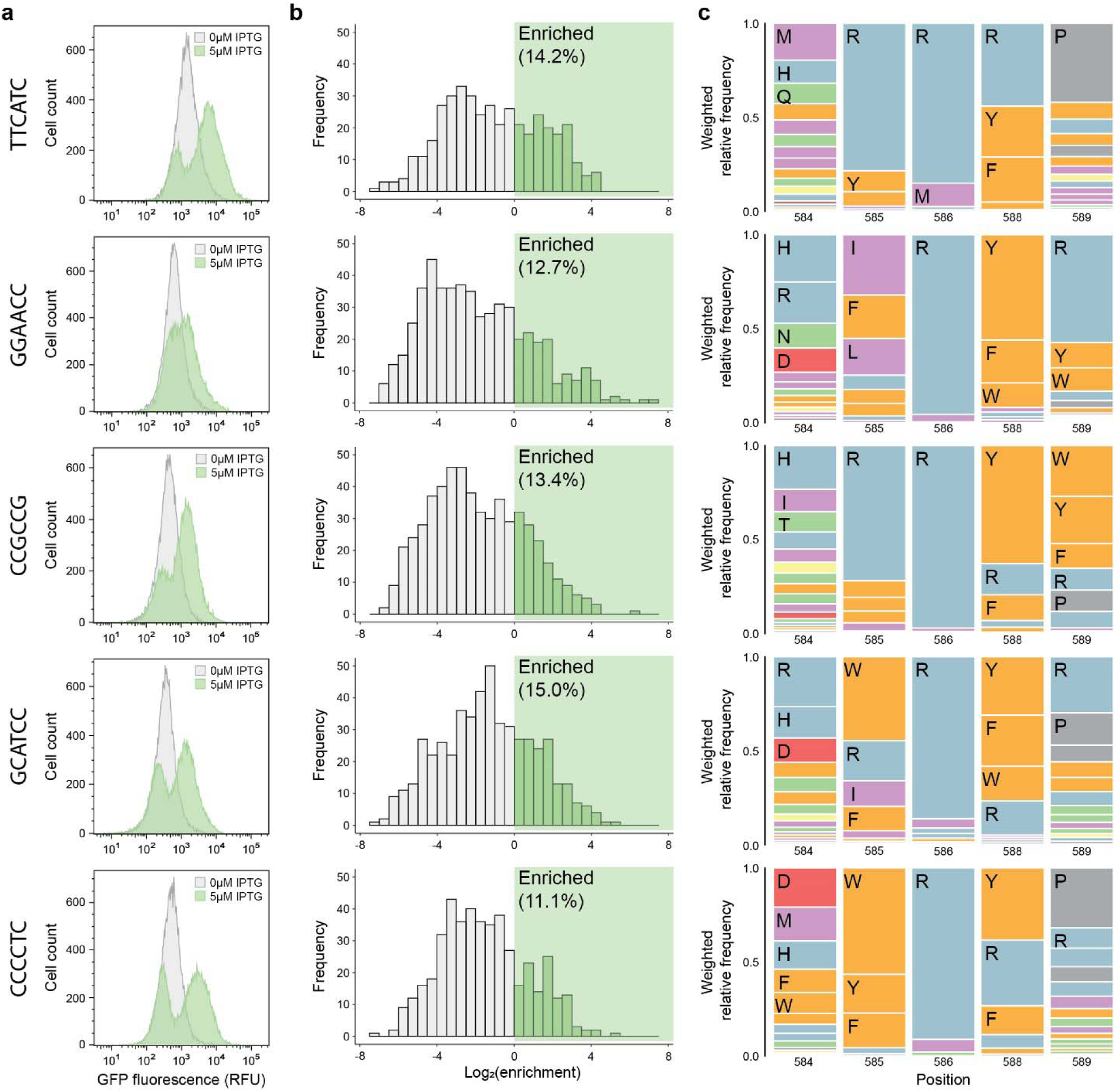
Selection and identification of successfully redesigned sigma-70 variants. (**a**), Flow cytometry fluorescence distributions of uninduced (gray) and IPTG induced (green) sigma-70 variant populations for each -35 target after FACS-based selection of functional redesigns. Transcriptionally active variants were enriched using two rounds of sequential GFP positive cell sorting. (**b**), Distributions of log-transformed enrichment scores of all characterized sigma-70 variants after selection. Deep sequencing was performed on the presorted and sorted libraries to compute enrichment ratios. (**c**), Sequence logos showing the weighted amino acid frequencies at each mutable position among functionally enriched sigma-70 variants. Amino acid identities are colored by chemical properties: polar amino acids (N, Q, S, T) shown in green, basic (H, K, R) blue, acidic (D, E) red, hydrophobic (A, I, L, M, V) purple, aromatic (F, W, Y) orange, and other (C, G, P) gray.

### Sequence profiles of redesigned sigma-70 variants are unique to each -35 target

Following deep sequencing, we evaluated the sigma-70 sequence profiles among enriched library variants of each -35 target. Given that the 5 promoter targets are separated by a median Hamming distance of 4, we expected the variable positions of sigma to undergo unique sequence selection (i.e., position-specific amino acid enrichment) to recognize each -35 target (**Supplementary Fig. 5**). While some similarities are shared across the sequence profiles for the five targets, each consensus sequence is unique (**Fig. 4c, Supplementary Fig. 4c**). Arginine was highly enriched at position 586, whereas both tyrosine and arginine were enriched at position 588 for all targets. In the WT sigma-70:DNA co-crystal structure (PDB: 4YLN) and Rosetta structural models, positions 586 and 588 interact primarily with the sugar-phosphate backbone of DNA (**Fig. 3c**). Thus, the enrichment of charged and/or polar residues at these positions is consistent with forming favorable non-specific DNA interactions. These structures also show base-specific interactions facilitated by position 585, which is where we observe several differences in enrichment across the promoter targets: arginine is favored by targets TTCATC and CCGCCG, bulky hydrophobic residues (I, F, and L) by GGAACC, aromatic residues (W, Y, and F) by CCCCTC and a wider range of residue types (W, R, I, and F) by GCATCC (**Fig. 4c**). Selection was less stringent at positions 584 and 589 of sigma-70 for all targets, as a broad range of amino acid identities were found among enriched variants. These positions often do not directly interact with the promoter in the Rosetta models. However, charged residues (R and H) were more enriched at positions 584 and 589 by targets GGAACC and GCATCC, which may electrostatically enhance DNA affinity. Additional interactions may be required to facilitate binding because the interactions by position 585 with the respective -35 targets are less specific. These results demonstrate that promoter specificity of sigma-70 is dictated by compounding differences in primary sequence that modulate both base-specific and general DNA affinity interactions.

To further validate that the targets were binding to the target -35 hexamer, we performed 5′ Rapid Amplification of cDNA Ends (5′ RACE) to determine the transcriptional start site (TSS) for two of our engineered Sigma 70 promoter pairs. In this technique, an RT-PCR is followed by the ligation of a 5′ adapter. Sanger sequencing, using the adapter and a gene-specific primer set, then allows for determination of the (TSS) by detecting the first nucleotide following the adapter sequence.

We conducted this experiment by overexpressing the wild-type (WT) Sigma70 with apFab71 Δ(-7), as well as examining RFHIMR with GGAACC-apFab71 Δ(-7) and FIQIRY with TTCACT-apFab71 Δ(-7). We found heterogeneous start sites for the WT Sigma70 apFab71 Δ(-7), with transcription occurring either at the canonical TSS or 9 bp upstream, although the Sanger mapping was poor (**Supplementary Fig. 6).** For the FIQIRY mutant, we observed a shift of 5 nucleotides downstream of the canonical TSS, and for the RFHIMR mutant, a shift of 7 nucleotides downstream. These shifts are similar to one of the observed TSS start sites for WT Sigma70, suggesting that either overexpression of the sigma factor or the modified -10 site leads to changes in the expected TSS. Despite these shifts, we found that the variant sigma factors initiate transcription in roughly the same region as the WT factor, indicating correct binding to the target -35 sites.

### Redesigned sigma-70’s exhibit varying levels of activity on the promoter targets

To validate the results of our high-throughput screen, we evaluated the activity of clonal isolates from our redesigned sigma-70 libraries. A total of 96 clones from the sorted libraries were randomly sampled, and from this set, we were able to identify redesigned sigma-70s with activity on each of the five promoter targets (**Fig. 5a, Supplementary Table 2**). To assess the performance of each sigma variant, we first measured the fluorescence of cells containing the target promoters upstream of GFP in the presence of endogenous WT sigma-70. Under this condition, the promoter variants yielded low fluorescence (<400 RFU OD^-1^) for the TTCATC, GGAACC, CCGCCG, GCTACC, and CCCCTC targets (**Fig. 5a**). In comparison, the activity of endogenous WT sigma-70 on the canonical -35 element (TTGACA) resulted in much higher fluorescence (10,200 ± 800 RFU OD^-1^) (**Supplementary Fig. 7**). Next, we measured the fluorescence of each clone on their respective -35 target sequence. Of the 96 colonies screened, 25-50% exhibited activity resulting in at least a four-fold increase in fluorescence over the endogenous WT sigma-70 baseline on the target promoter (**Supplementary Table 2**). The proportion of ‘successful’ redesigns in the sorted libraries is consistent with the flow cytometry profiles (**Fig. 4a**). The activities of the top three performing sigma-70 redesigns for each target varied across the targets. The best performing design had 77% of the activity of endogenous WT sigma-70 on the canonical -35 element (**Fig. 5a**). For the target TTCATC, which is closest in Hamming distance to the canonical -35 sequence (**Fig. 2d**), we found the three highest performing redesigned sigma-70s (CRRVY, FIQRY, and FWCRY). However, performance does not scale with Hamming distance, suggesting a more complex relationship exists between a given - 35 target and the success of computationally redesigned sigma-70s.

**Fig. 5.**
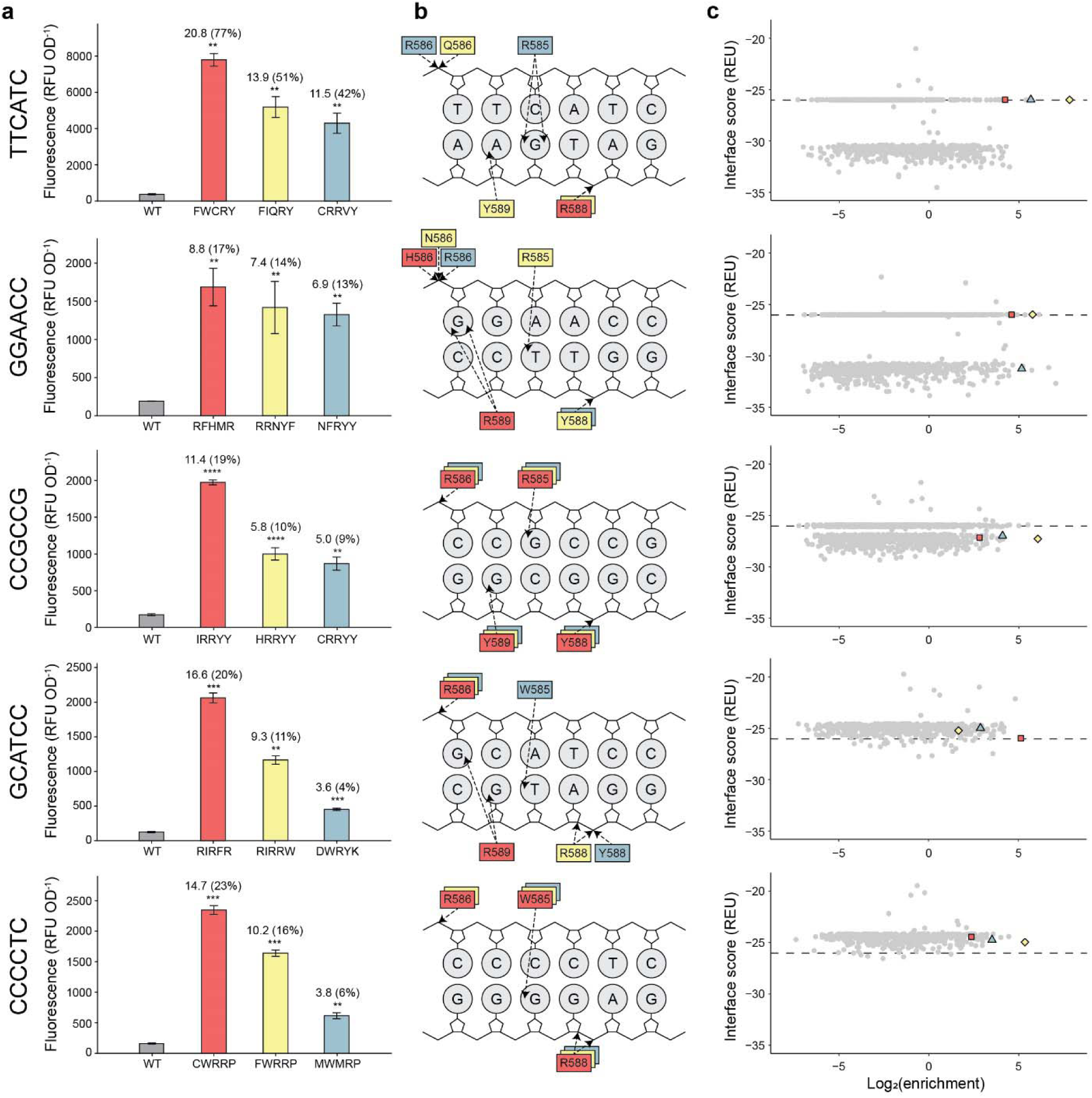
Clonal validation of redesigned sigma-70 variants. (**a**), Normalized fluorescence of WT sigma-70 (gray) and the top three performing variants (red, yellow, or blue) for each promoter target. Clones were selected from the sorted libraries and assayed in a 96-well fluorescence plate reader. Error bars denote the standard deviation (**P ≤ 0.01, ***P ≤ 0.001) of replicates (n≥3) and computed fold-changes are relative to the fluorescence resulting from endogenous WT sigma-70 with each target. Percentages of activity are relative to the activity of endogenous WT sigma-70 with the promoter containing a canonical -35 element (**Supplementary Fig. 6**). RFU, relative fluorescence units. OD, optical density at 600nm. (**b**), Cartoon schematic showing H-bonds formed between each sigma-70 variant and the cognate -35 DNA target in the Rosetta structural models. (**c**), Rosetta interface scores and enrichment scores from the high-throughput screen of all tested sigma-70 variants. The top three performing variants are indicated in red (square), yellow (circle), and blue (triangle).

### Rosetta structural models reveal potential mechanisms of -35 target recognition

In addition to the low transcriptional activation by WT sigma-70, the five -35 targets were selected due to their sequence deviation (i.e. orthogonality) from the canonical -35 and each other. The utility of engineered sigma factors is dependent on minimizing crosstalk between various sigma factor-target pairs. By targeting diverse -35 sequences with our redesign of sigma-70, we sought to generate novel and specific protein-DNA interactions for each sigma factor-target pair. Evaluating Rosetta structural models generated for the highest performing sigma-70 redesigns, we observe several differences between the hydrogen bonding patterns of our redesigned sigma-70 variants with their cognate -35 targets (**Fig. 5b).** Some interactions were largely conserved across target sequences, such as the h-bonds between residues 586 and 588 of sigma and the sugar-phosphate backbone of nucleotides -35 and -31, despite the diverse polar and charged residues employed to maintain these interactions. In contrast to the other -35 targets, the hydrogen bonding patterns of the top three performing sigma-70 redesigns for targets CCGCCG and CCCCTC are highly converged. The base-specific interactions between CCGCCG and variants IRRYY, HRRYY, and CRRYY occur between R585:Gua-33 and Y589:Gua-34. For CCCCTC, a single base-specific interaction, W585:Gua-33, appears sufficient for recognition by sigma-70 variants CWRRP, FWRRP, and MWMRP, perhaps necessitating the additional non-specific h-bond between R586 and the sugar-phosphate backbone. However, the number of h-bonds present in these protein-DNA structural models ranges from one to four, which highlights the importance of other types of molecular interactions (van der Waal’s, dipole-dipole) for promoter recognition. Though computationally derived, these models show how unique sets of interactions may facilitate -35 target recognition by our successfully redesigned sigma-70 variants.

To understand the relationship between the Rosetta interface scores and the performance of each sigma-70 redesign, we compared these energies to the enrichment scores from our high-throughput screen (**Fig. 5c**). While the computational screening approach was successful at generating functional variants and significantly reduced the sequence space search from 3.2 million (20^5^) combinations to a manageable 1000-member test set, the highest affinity redesigns (i.e. lowest interface scores) were not among the best performing. This underscores the necessity of our high-throughput screening platform using florescent activated cell sorting (FACS) and deep sequencing to isolate functional redesigns. The top three performing sigma-70 variants for each -35 target had Rosetta interface scores comparable to that of WT sigma-70 on the canonical -35 element (-26.0 REU), suggesting that an ‘optimal binding affinity’ exists for DNA recognition and transcription initiation by sigma-70. An optimized affinity may facilitate proper association with the promoter and allow dissociation of sigma-70 to initiate transcription and elongation.

## DISCUSSION

We devised a high throughput *in silico* modeling and *in vivo* screening workflow to redesign promoter specificity of the *E. coli* “housekeeping” regulator of transcription, sigma-70. Using this workflow, we identified multiple sigma-70 variants that activate transcription on 5 target promoters containing diverse - 35 elements: TTCATC, GGAACC, CCGCCG, GCTACC, and CCCCTC. These results demonstrate that the promoter specificity of an essential primary transcription regulator can be rationally redesigned. Although the process of transcription initiation (i.e. promoter recognition and DNA melting) by sigma factors is complex, the partitioned functional domains of sigma factors enabled key interactions between the domain 4 recognition helix and the -35 DNA element to be redesigned without perturbing other functions. Our success across all five target sequences demonstrates the extant at which promoter specificity can be altered and the generalizability of our approach. Given that the Hamming distance separation ranges from three to six (of 6 nucleotide positions) between these novel -35 elements and the canonical -35 element, TTGACA, our approach also enables the design of sigma factors with maximal orthogonality from natural sigma factors and thereby, greater potential to minimize crosstalk between synthetic and host expression systems. In contrast, sigma factors generated in previous works using domain swaps of alternative sigma factors or introduced from non-host family bacteria, have been limited in their orthogonality and host compatibility.

One advantage to the engineered sigma-70 variants presented is that they integrate seamlessly into the *E. coli* host, as they utilize the same endogenous transcriptional machinery. While the level of activation varied from 3-fold to 20-fold across our top performing redesigns, we show that this activity was also engineered *de novo*, as the native sigma-70 exhibited little to no activity on each of the orthogonal promoters. Using structural models of these successful redesigns in complex with their cognate promoters, we observe diverse combinations of interactions that facilitate promoter recognition. This diversity extends even among successful redesigns for the same promoter target, demonstrating the utility of computational tools to navigate the energetic landscape and simplify the sequence space to be experimentally tested.

The transcription-level regulatory complexity of a cell is mediated by the concerted efforts of global and local regulators. While many existing synthetic expression systems are reliant solely on local regulators (i.e. natural or synthetic transcription factors) to control a sparse set of genes, they lack the complexity and capabilities of native systems. Only by incorporating global regulatory components, such as engineered sigma factors, can we expand the utility and complexity of synthetic genetic circuits. However, building or isolating an extensive repertoire of global regulatory components is challenging because they must be (1) orthogonal to host systems to minimize crosstalk between the synthetic and native expression networks, (2) display dynamic control by activating gene expression in response to simple input signals, and (3) be highly customizable and modular, such that they can be easily integrated into established expression systems. Here, we developed and validated a generalizable approach to engineer sigma-70 variants that satisfy all these requirements and can be similarly applied to any host sigma factor to expand the set of available global regulatory components. Future work could pair these engineered sigma-70 factors with existing local regulators of transcription in a biosynthetic expression system.

## ACKNOWLEDGEMENTS

This work was partly supported by the NIH Director’s New Innovator’s Award (DP2GM132682-01 to S.R.), partly by the Great Lakes Bioenergy Research Center, U. S. Department of Energy, Office of Science, Office of Biological and Environmental Research (Award Number DE-SC0018409) and partly by the NIGMS Biotechnology Training Program (T32GM135066 to J.O.S). The content is solely the responsibility of the authors and does not necessarily represent the official views of the NIH, Department of Defense, Department of Energy, or other federal agencies.

## AUTHOR CONTRIBUTIONS

X.L., A.T.M., T.G., and S.R. conceptualized the project. X.L. devised the cell-based screening system, cloned the promoter library, and performed sorting and NGS experiments. A.T.M. performed the *in silico* calculations and designed the sigma variant libraries. T.G. performed clonal screens and identified the top performing sigma variants for each promoter target. J.O.S. performed supplementary experiments. X.L., A.T.M., T.G, J.O.S., and S.R. drafted the manuscript. A.T.M., X.L., T.G., and J.O.S created the figures. B.S .and R.L. performed 5 prime RACE experiments. S.R. supervised the work. All authors edited and approved the final manuscript.

## Notes

### Competing Interest Statement

The authors have declared no competing interest.

### Summary of Updates

We have added new experimental determining the transcriptional start site of the designed sigma proteins.

