## Supplementary data for "Computation-guided redesign of promoter specificity of a bacterial RNA polymerase"

**SUPPLEMENTARY INFORMATION**

**Supplementary Table 1**. **Normalized fluorescence measurements of clones containing randomized -35 sequences.** GFP fluorescence and OD_600nm_ measurements were collected with a 96-well plate reader. Hamming distances for each sequence are relative to the canonical -35 element. The five selected low activity -35 targets are highlighted in green.

| -35 sequence | Normalized fluorescence  (RFU OD^-1^) | Hamming distance from TTGACA |
| --- | --- | --- |
| ATGACT | 316 | 2 |
| AAGCCA | 426 | 3 |
| TTATCC | 488 | 3 |
| TTCATC | 320 | 3 |
| TTCTCC | 492 | 3 |
| AGGATG | 627 | 4 |
| ATACCC | 340 | 4 |
| ATATCC | 381 | 4 |
| ATCTCT | 420 | 4 |
| CCGCCC | 445 | 4 |
| CCGCCG | 491 | 4 |
| CGGCCC | 474 | 4 |
| CTCGCC | 426 | 4 |
| CTGCTC | 452 | 4 |
| CTTTCT | 436 | 4 |
| GCGAGT | 404 | 4 |
| GCGCCT | 548 | 4 |
| GCGGCC | 413 | 4 |
| GGAACC | 585 | 4 |
| GTCGCC | 439 | 4 |
| TCTCCC | 483 | 4 |
| TTAGTG | 494 | 4 |
| TTCTTT | 514 | 4 |
| AAACCC | 601 | 5 |
| AAAGTA | 497 | 5 |
| AACGCC | 475 | 5 |
| AAGTGT | 341 | 5 |
| ACGGTT | 434 | 5 |
| AGACCC | 376 | 5 |
| AGAGCC | 431 | 5 |
| AGTTCC | 478 | 5 |
| ATTTTT | 479 | 5 |
| CCCTCC | 416 | 5 |
| CCTTCT | 487 | 5 |
| CGAGCC | 413 | 5 |
| CGCCCG | 523 | 5 |
| CGCGCC | 396 | 5 |
| CGGCTC | 527 | 5 |
| CGTCCT | 533 | 5 |
| CGTGCC | 394 | 5 |
| CGTTCT | 489 | 5 |
| CTCCTT | 459 | 5 |
| GACTCT | 422 | 5 |
| GATCCC | 421 | 5 |
| GATGCT | 397 | 5 |
| GATTCC | 436 | 5 |
| GCATCC | 454 | 5 |
| GCATCT | 491 | 5 |
| GCTTCT | 560 | 5 |
| GGCCCC | 310 | 5 |
| GTCTTT | 510 | 5 |
| GTTTTT | 473 | 5 |
| TACTTT | 311 | 5 |
| TATTTT | 376 | 5 |
| TGCGTT | 327 | 5 |
| TGCTTC | 572 | 5 |
| AATTTT | 421 | 6 |
| ACATTT | 402 | 6 |
| CACTTT | 442 | 6 |
| CATTGC | 487 | 6 |
| CCCCTC | 554 | 6 |
| CCTTTC | 521 | 6 |
| CCTTTT | 314 | 6 |
| CGCTTT | 328 | 6 |
| CGTCTT | 509 | 6 |
| CGTTTC | 409 | 6 |
| CGTTTT | 401 | 6 |
| GAATGT | 316 | 6 |
| GACTTT | 354 | 6 |
| GATCTT | 457 | 6 |
| GATTTT | 363 | 6 |
| GCCTTT | 349 | 6 |
| GCTTTT | 400 | 6 |
| GGCTTT | 413 | 6 |

**Supplementary Table 2. Plasmids and DNA Parts Utilized in this Study.**

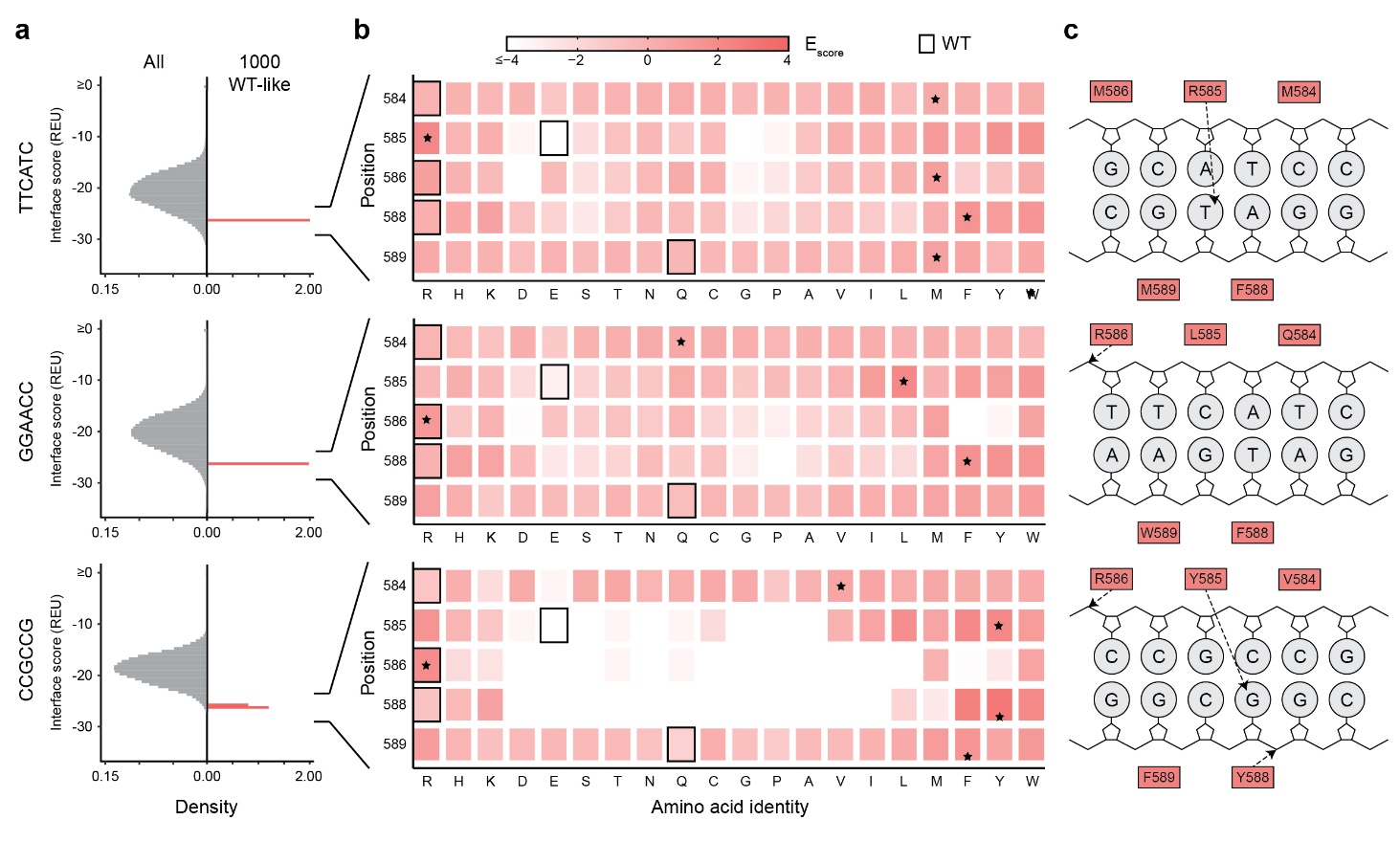

**Supplementary Fig. 1. Computation-based selection of redesigned sigma-70 variants for three promoter targets.** (**a**), Interface scores of all (gray) or 1000 most WT-like (Interface score = -26REU, red) sigma-70 variants in complex with each -35 DNA target. WT-like sets were not created for promoter targets CCCCTC and GCATCC because they were redundant with the lowest energy sets. All single, double, triple, and quadruple combinatorial variants of sigma-70 positions 584, 585, 586, 588, and 589 were modeled using Rosetta. (**b**), Position-specific amino acid enrichment scores (red gradient) among selected wt-like scoring sigma-70 variants. WT identity (boxed outline) and most enriched amino acid (*) at each mutable position. (**c**), Cartoon schematic showing H-bonds formed between each enriched sigma-70 consensus sequence and -35 DNA target in the Rosetta structural models.

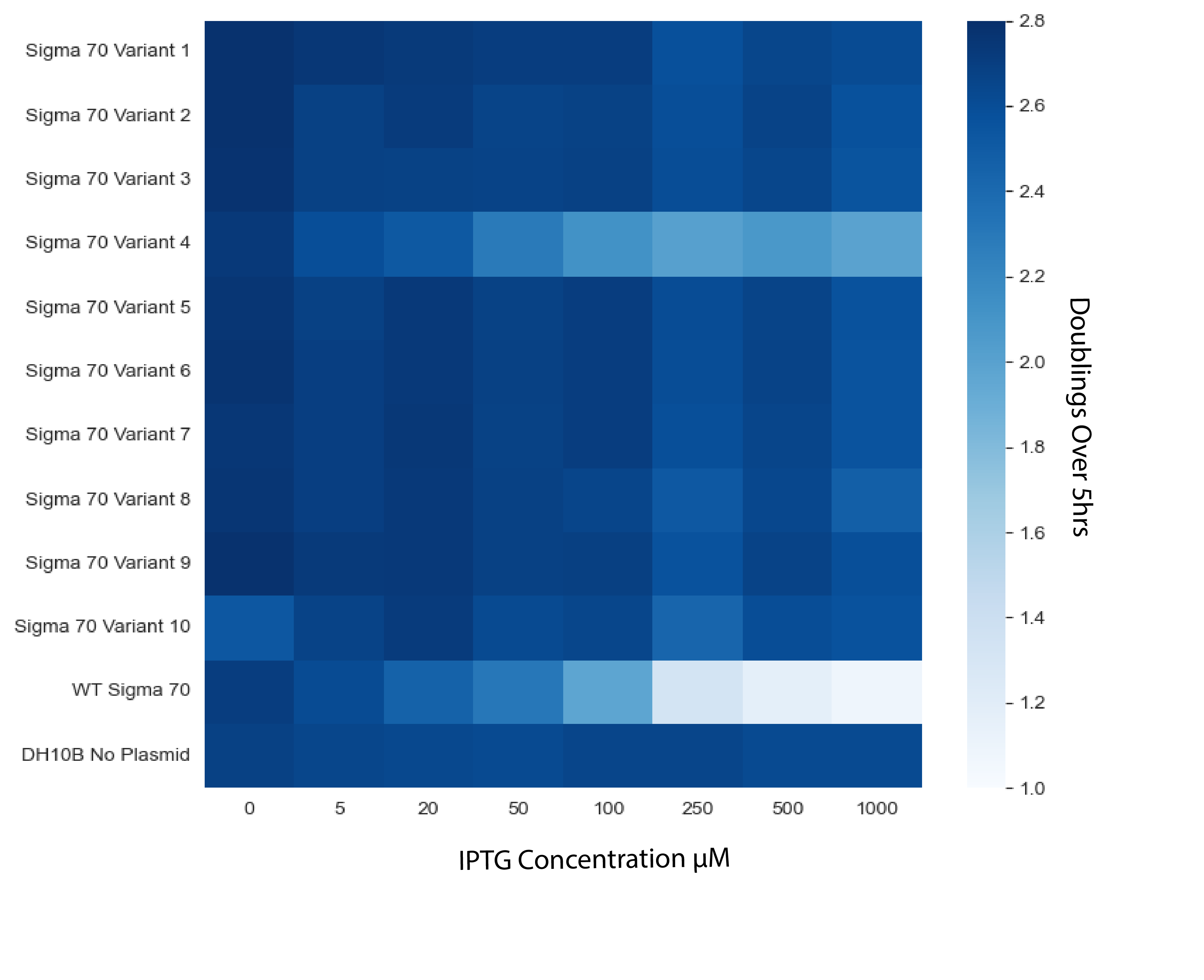

**Supplementary Fig. 2. Sigma-70** **expression induced toxicity**. Each row corresponds to a different random sigma-70 variant picked from the library. Each column represents a different IPTG concentration. Color coding represents cell doublings measured by OD_600_nm after 5 hours of induction. Dark represents high cell density while white represents low cell density.

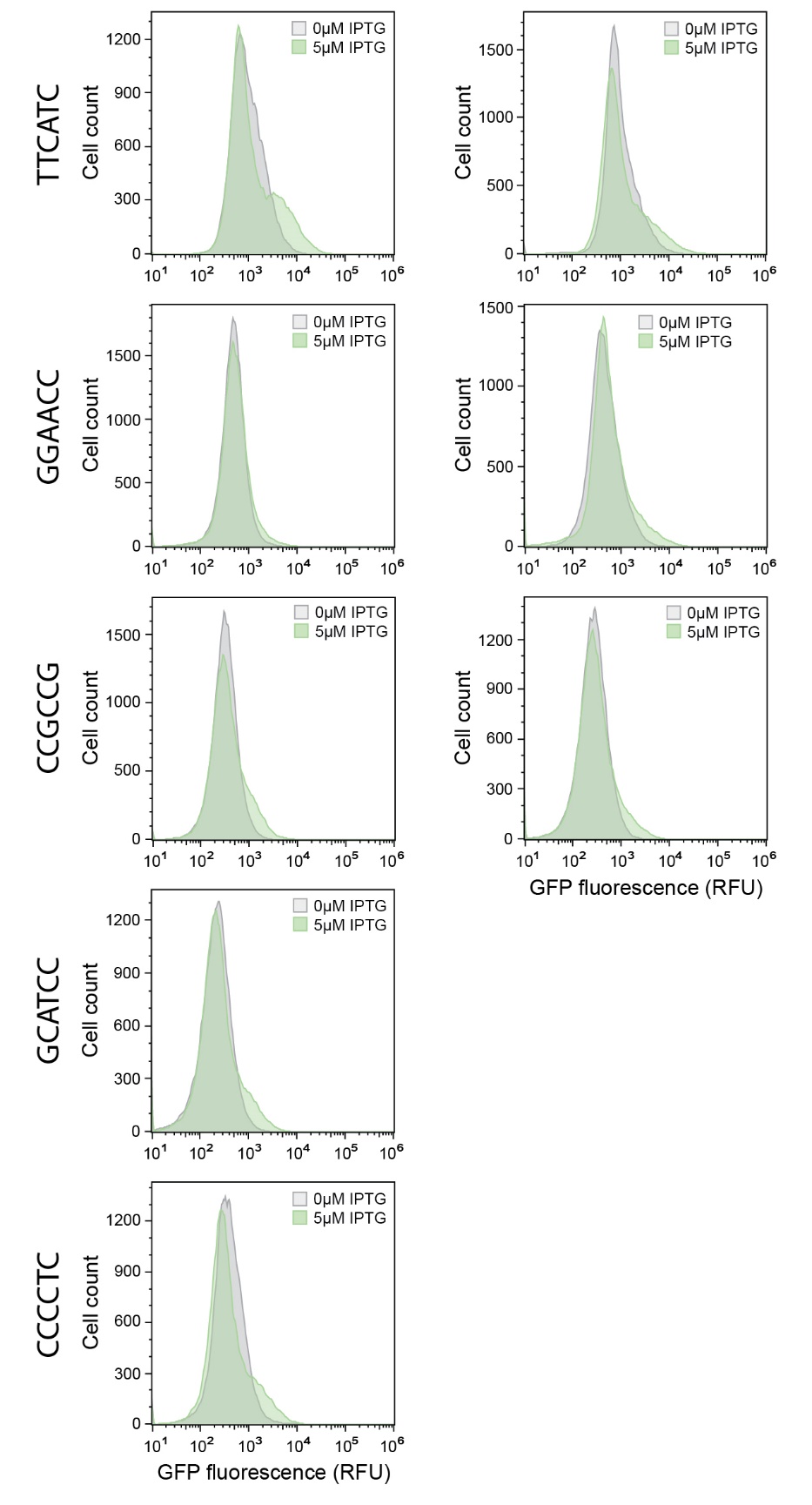

**Fig. 3. Selection of successfully redesigned sigma-70 variants.** Flow cytometry fluorescence distributions of uninduced (gray) and IPTG induced (green) sigma-70 variant populations for each -35 target prior to FACS-based selection of functional redesigns. Transcriptionally active variants were enriched by sorting the GFP positive subpopulation of the lowest energy (left) and WT-like (right) libraries of each target.

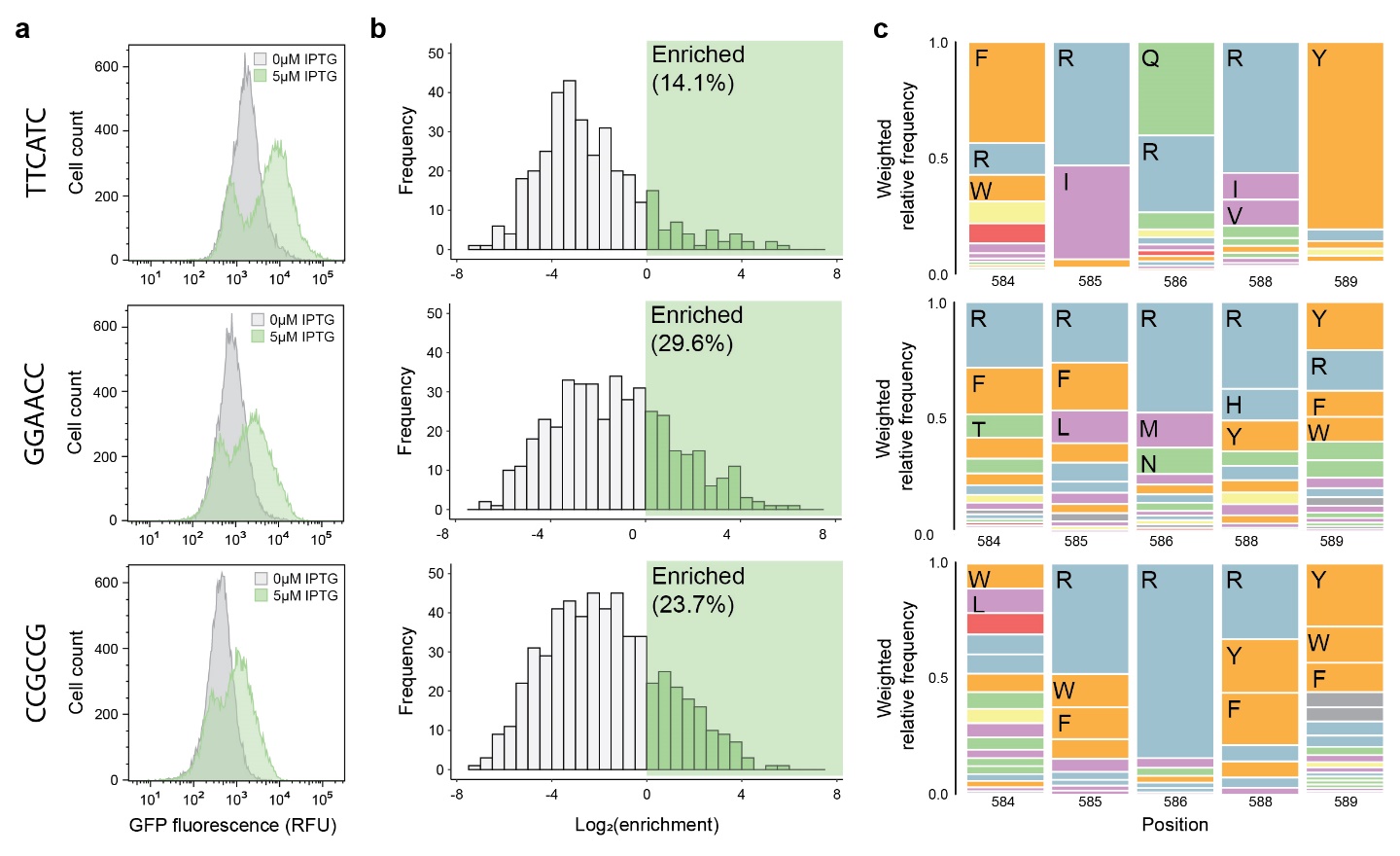

**Supplementary Fig. 4. Selection and identification of successfully redesigned sigma-70 variants.** (**a**), Flow cytometry fluorescence distributions of uninduced (gray) and IPTG induced (green) sigma-70 variant populations for three -35 targets after FACS-based selection of functional redesigns. Transcriptionally active variants from the WT-like libraries were enriched using two rounds of sequential GFP positive cell sorting. (**b**), Distributions of log-transformed enrichment scores of all characterized sigma-70 variants after selection. Deep sequencing was performed on the presorted and sorted libraries to compute enrichment ratios. (**c**), Sequence logos showing the weighted amino acid frequencies at each mutable position among functionally enriched sigma-70 variants. Amino acid identities are colored by chemical properties: polar amino acids (N, Q, S, T) shown in green, basic (H, K, R) blue, acidic (D, E) red, hydrophobic (A, I, L, M, V) purple, aromatic (F, W, Y) orange, and other (C, G, P) gray.

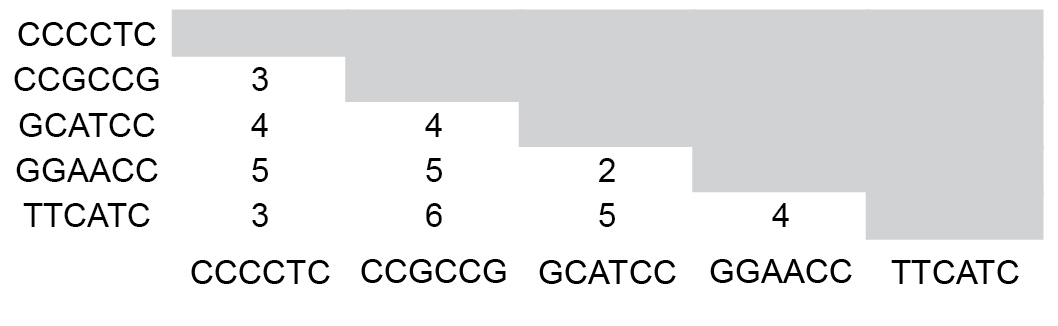

**Supplementary Fig. 5. Hamming distances between each of the five selected -35 targets.**

**Supplementary Table 2. Clonal screens of sigma variants after FACS-based selection of high activity variants.** GFP fluorescence and OD_600nm_ measurements were collected with a 96-well plate reader. 96 clones were tested for each -35 target. Normalized GFP fluorescence was compared to WT sigma on each target to compute fold-improvement.

| -35 target | Total number of  clones screened | Number of clones with  fold-improvement ≥4 |
| --- | --- | --- |
| GGAACC | 96 | 50 |
| CCCCTC | 96 | 21 |
| TTCATC | 96 | 51 |
| GCATCC | 96 | 23 |
| CCGCCG | 96 | 35 |

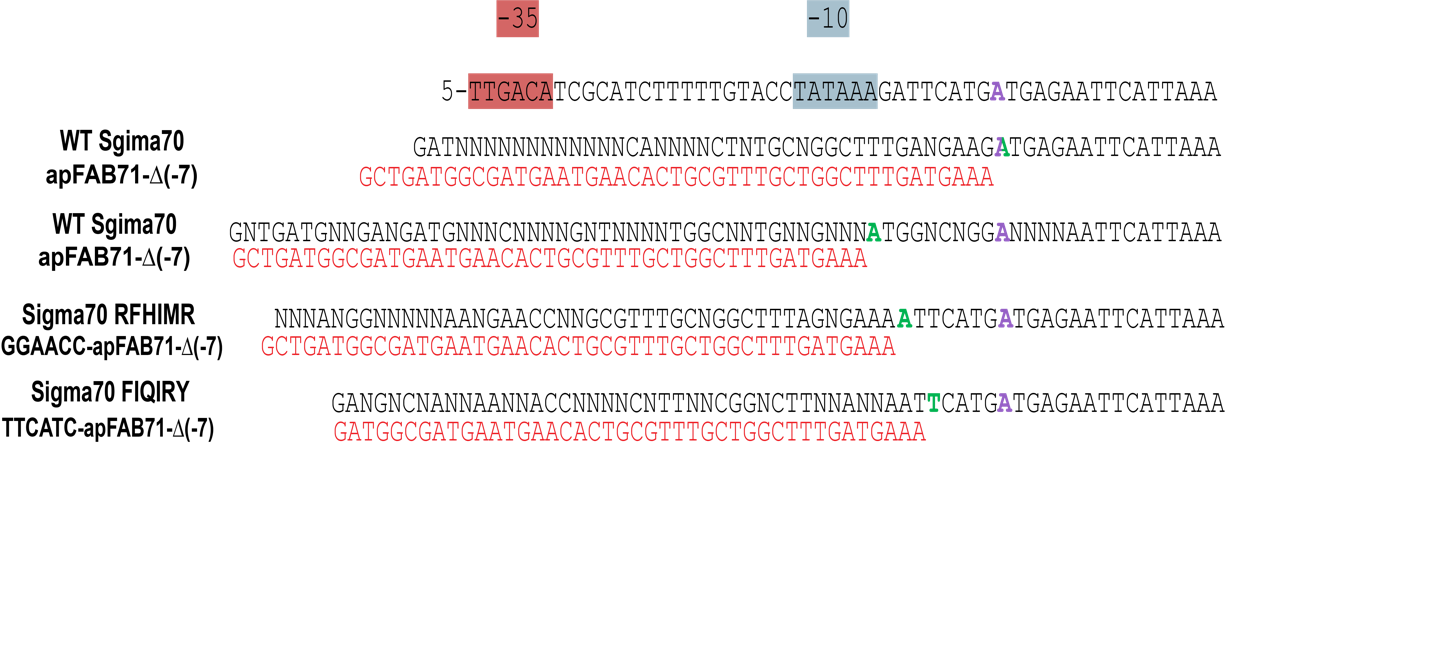

**Supplementary Fig, 6. 5**′ **RACE Characterization of RFHIMR and FIQIRY Variants on their Target Promoters.** Alignment of the sequencing results to apFAB71(∆-7) (top sequence) and the 5′ RACE adapter (red sequence). The canonical transcriptional start site is in purple while the experimentally determined transcriptional start sites are in green. Overexpressed WT Sigma70 demonstrated two TSS, one at the canonical location and another -9 bp away, The Sigma 70 variants RFHMIR and FIQIRY had a -7 and -5 nucleotide shift respectively.

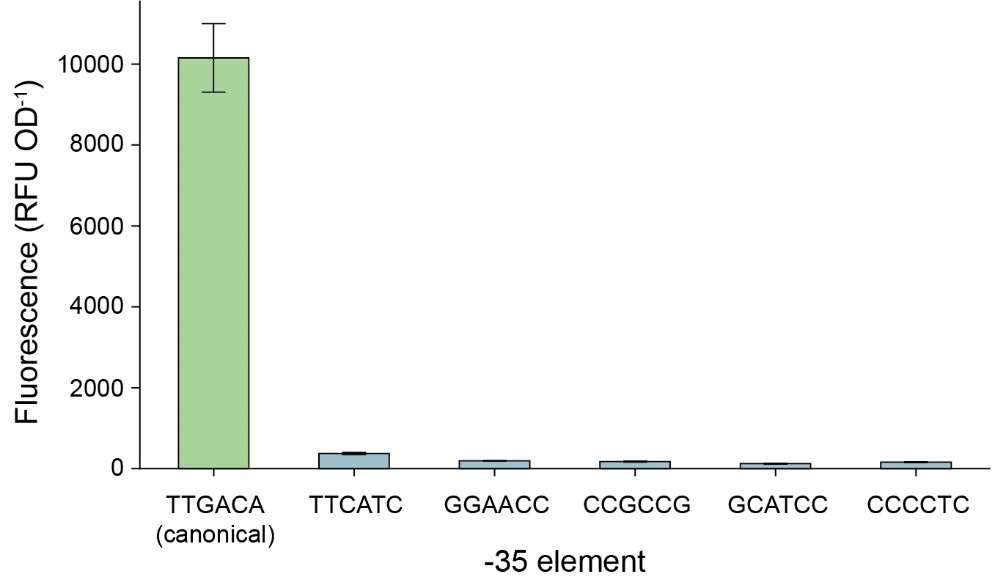

**Supplementary Fig. 7. Activity of endogenous WT sigma-70 on the canonical and target -35 elements.** GFP fluorescence and OD_600nm_ measurements (n ≥ 3) were collected with a 96-well plate reader.

**Supplementary Table 3. Select plasmid sequences and parts utilized in this experiment.** For target promoters the -35 and -10 sites are underlined.

| Plasmid/Part | Purpose | Sequence |
| --- | --- | --- |
| PXL-9 | Reporter Plasmid | CGTTCGGCTGCGGCGAGCGGTATCAGCTCACTCAAAGGCGGTAATACGGTTATCCACAGAATCAGGGGATAACGCAGGAAAGAACATGTGAGCAAAAGGCCAGCAAAAGGCCAGGAACCGTAAAAAGGCCGCGTTGCTGGCGTTTTTCCATAGGCTCCGCCCCCCTGACGAGCATCACAAAAATCGACGCTCAAGTCAGAGGTGGCGAAACCCGACAGGACTATAAAGATACCAGGCGTTTCCCCCTGGAAGCTCCCTCGTGCGCTCTCCTGTTCCGACCCTGCCGCTTACCGGATACCTGTCCGCCTTTCTCCCTTCGGGAAGCGTGGCGCTTTCTCATAGCTCACGCTGTAGGTATCTCAGTTCGGTGTAGGTCGTTCGCTCCAAGCTGGGCTGTGTGCACGAACCCCCCGTTCAGCCCGACCGCTGCGCCTTATCCGGTAACTATCGTCTTGAGTCCAACCCGGTAAGACACGACTTATCGCCACTGGCAGCAGCCACTGGTAACAGGATTAGCAGAGCGAGGTATGTAGGCGGTGCTACAGAGTTCTTGAAGTGGTGGCCTAACTACGGCTACACTAGAAGGACAGTATTTGGTATCTGCGCTCTGCTGAAGCCAGTTACCTTCGGAAAAAGAGTTGGTAGCTCTTGATCCGGCAAACAAACCACCGCTGGTAGCGGTGGTTTTTTTGTTTGCAAGCAGCAGATTACGCGCAGAAAAAAAGGATCTCAAGAAGATCCTTTGATCTTTTCTACGGGGTCTGACGCTCAGTGGAACGAAAACTCACGTTAAGGGATTTTGGTCATGACTAGTGCTTGGATTCTCACCAATAAAAAACGCCCGGCGGCAACCGAGCGTTCTGAACAAATCCAGATGGAGTTCTGAGGTCATTACTGGATCTATCAACAGGAGTCCAAGCGAGCTCTCGAACCCCAGAGTCCCGCTCAGAAGAACTCGTCAAGAAGGCGATAGAAGGCGATGCGCTGCGAATCGGGAGCGGCGATACCGTAAAGCACGAGGAAGCGGTCAGCCCATTCGCCGCCAAGCTCTTCAGCAATATCACGGGTAGCCAACGCTATGTCCTGATAGCGGTCCGCCACACCCAGCCGGCCACAGTCGATGAATCCAGAAAAGCGGCCATTTTCCACCATGATATTCGGCAAGCAGGCATCGCCATGGGTCACGACGAGATCCTCGCCGTCGGGCATGCGCGCCTTGAGCCTGGCGAACAGTTCGGCTGGCGCGAGCCCCTGATGCTCTTCGTCCAGATCATCCTGATCGACAAGACCGGCTTCCATCCGAGTACGTGCTCGCTCGATGCGATGTTTCGCTTGGTGGTCGAATGGGCAGGTAGCCGGATCAAGCGTATGCAGCCGCCGCATTGCATCAGCCATGATGGATACTTTCTCGGCAGGAGCAAGGTGAGATGACAGGAGATCCTGCCCCGGCACTTCGCCCAATAGCAGCCAGTCCCTTCCCGCTTCAGTGACAACGTCGAGCACAGCTGCGCAAGGAACGCCCGTCGTGGCCAGCCACGATAGCCGCGCTGCCTCGTCCTGCAGTTCATTCAGGGCACCGGACAGGTCGGTCTTGACAAAAAGAACCGGGCGCCCCTGCGCTGACAGCCGGAACACGGCGGCATCAGAGCAGCCGATTGTCTGTTGTGCCCAGTCATAGCCGAATAGCCTCTCCACCCAAGCGGCCGGAGAACCTGCGTGCAATCCATCTTGTTCAATCATGCGAAACGATCCTCATCCTGTCTCTTGATCAGATCTTGATCCCCTGCGCCATCAGATCCTTGGCGGCAAGAAAGCCATCCAGTTTACTTTGCAGGGCTTCCCAACCTTACCAGAGGGCGCCCCAGCTGGCAATTCCGACGTCTAAGAAACCATTATTATCATGACATTAACCTATAAAAATAGGCGTATCACGAGGCCCTTTCGTCTTCACCTCGAGCGGCCGCAAAAGGAAAAGATCCGGCAAACAAACCACCGTTGGTAGCGGTGGTTTTTTTGTTTGGATCGACAATCTTCGTAAGCGTCATCAATAAGCGTAAAAAAACCGGGCAATGCCCGGTTTTTTAATGAGAAATTTTACCTGTCGTAGCCGCCACCATCCGGCAAAGAAGCATACAAGGCTTTTGGCTTATAGCTACGTAGCGCATTGCGTCGCAGCACAATCCCGGCACCGATCAAGTCTTCGCGATGATTATTATTATTTGTACAGCTCATCCATGCCACCGGTAGAATCTGATTGACATCGCATCTTTTTGTACCTATAATAGATTCATGATGAGAATTCATTAAAGAGGAGAAAGGTCATATGCGTAAAGGCGAAGAGCTGTTCACTGGTTTCGTCACTATTCTGGTGGAACTGGATGGTGATGTCAACGGTCATAAGTTTTCCGTGCGTGGCGAGGGTGAAGGTGACGCAACTAATGGTAAACTGACGCTGAAGTTCATCTGTACTACTGGTAAACTGCCGGTACCTTGGCCGACTCTGGTAACGACGCTGACTTATGGTGTTCAGTGCTTTGCTCGTTATCCGGACCACATGAAGCAGCATGACTTCTTCAAGTCCGCCATGCCGGAAGGCTATGTGCAGGAACGCACGATTTCCTTTAAGGATGACGGCACGTACAAAACGCGTGCGGAAGTGAAATTTGAAGGCGATACCCTGGTAAACCGCATTGAGCTGAAAGGCATTGACTTTAAAGAAGACGGCAATATCCTGGGCCATAAGCTGGAATACAATTTTAACAGCCACAATGTTTACATCACCGCCGATAAACAAAAAAATGGCATTAAAGCGAATTTTAAAATTCGCCACAACGTGGAGGATGGCAGCGTGCAGCTGGCTGATCACTACCAGCAAAACACTCCAATCGGTGATGGTCCTGTTCTGCTGCCAGACAATCACTATCTGAGCACGCAAAGCGTTCTGTCTAAAGATCCGAACGAGAAACGCGATCACATGGTTCTGCTGGAGTTCGTAACCGCAGCGGGCATCACGCATGGTATGGATGAACTGTACAAATAATAATGCAGGTCGTCTCGGATCGAGAAGGACACGGTTAATACTAGGCCTGCTGGCTGGTAATCGCCAGCAGGCCTTTTTATTTGGGGGAGAGGGAAGTCATGAAAAAACTAACCTTTGAAATTCGATCTCCACCACATCAGCTCTGAAGCAACGTAAAAAAACCCGCCCCGGCGGGTTTTTTTATACCCGTAGTATCCCCACTTATCTACAATAGCTGTCCTTAATTAATCTAGA |
| SC101_LacI_Sigma_70 | Sigma Factor Expression Plasmid | AATTGGTCTAGGTGATTTTAATCACTATACCAATTGAGATGGGCTAGTCAATGATAATTACATGTCCTTTTCCTTTGAGTTGTGGGTATCTGTAAATTCTGCTAGACCTTTGCTGGAAAACTTGTAAATTCTGCTAGACCCTCTGTAAATTCCGCTAGACCTTTGTGTGTTTTTTTTGTTTATATTCAAGTGGTTATAATTTATAGAATAAAGAAAGAATAAAAAAAGATAAAAAGAATAGATCCCAGCCCTGTGTATAACTCACTACTTTAGTCAGTTCCGCAGTATTACAAAAGGATGTCGCAAACGCTGTTTGCTCCTCTACAAAACAGACCTTAAAACCCTAAAGGCTTAAGTAGCACCCTCGCAAGCTCGGGCAAATCGCTGAATATTCCTTTTGTCTCCGACCATCAGGCACCTGAGTCGCTGTCTTTTTCGTGACATTCAGTTCGCTGCGCTCACGGCTCTGGCAGTGAATGGGGGTAAATGGCACTACAGGCGCCTTTTATGGATTCATGCAAGGAAACTACCCATAATACAAGAAAAGCCCGTCACGGGCTTCTCAGGGCGTTTTATGGCGGGTCTGCTATGTGGTGCTATCTGACTTTTTGCTGTTCAGCAGTTCCTGCCCTCTGATTTTCCAGTCTGACCACTTCGGATTATCCCGTGACAGGTCATTCAGACTGGCTAATGCACCCAGTAAGGCAGCGGTATCATCAACAGGCTTACCCGTCTTACTGTCCCTAGTGCTTGGATTCTCACCAATAAAAAACGCCCGGCGGCAACCGAGCGTTCTGAACAAATCCAGATGGAGTTCTGAGGTCATTACTGGATCTATCAACAGGAGTCCAAGCGAGCTCGTAAACTTGGTCTGACAGAATGCAGGAGTCGCATAAGGGAGAGCGTCGAGATCCCGGACACCATCGAATGGCGCAAAACCTTTCGCGGTATGGCATGATAGCGCCCGGAAGAGAGTCAATTCAGGGTGGTGAATGTGAAACCAGTAACGTTATACGATGTCGCAGAGTATGCCGGTGTCTCTTATCAGACCGTTTCCCGCGTGGTGAACCAGGCCAGCCACGTTTCTGCGAAAACGCGGGAAAAAGTGGAAGCGGCGATGGCGGAGCTGAATTACATTCCCAACCGCGTGGCACAACAACTGGCGGGCAAACAGTCGTTGCTGATTGGCGTTGCCACCTCCAGTCTGGCCCTGCACGCGCCGTCGCAAATTGTCGCGGCGATTAAATCTCGCGCCGATCAACTGGGTGCCAGCGTGGTGGTGTCGATGGTAGAACGAAGCGGCGTCGAAGCCTGTAAAGCGGCGGTGCACAATCTTCTCGCGCAACGCGTCAGTGGGCTGATCATTAACTATCCGCTGGATGACCAGGATGCCATTGCTGTGGAAGCTGCCTGCACTAATGTTCCGGCGTTATTTCTTGATGTCTCTGACCAGACACCCATCAACAGTATTATTTTCTCCCATGAAGACGGTACGCGACTGGGCGTGGAGCATCTGGTCGCATTGGGTCACCAGCAAATCGCGCTGTTAGCGGGCCCATTAAGTTCTGTCTCGGCGCGTCTGCGTCTGGCTGGCTGGCATAAATATCTCACTCGCAATCAAATTCAGCCGATAGCGGAACGGGAAGGCGACTGGAGTGCCATGTCCGGTTTTCAACAAACCATGCAAATGCTGAATGAGGGCATCGTTCCCACTGCGATGCTGGTTGCCAACGATCAGATGGCGCTGGGCGCAATGCGCGCCATTACCGAGTCCGGGCTGCGCGTTGGTGCGGATATCTCGGTAGTGGGATACGACGATACCGAAGACAGCTCATGTTATATCCCGCCGTTAACCACCATCAAACAGGATTTTCGCCTGCTGGGGCAAACCAGCGTGGACCGCTTGCTGCAACTCTCTCAGGGCCAGGCGGTGAAGGGCAATCAGCTGTTGCCCGTCTCACTGGTGAAAAGAAAAACCACCCTGGCGCCCAATACGCAAACCGCCTCTCCCCGCGCGTTGGCCGATTCATTAATGCAGCTGGCACGACAGGTTTCCCGACTGGAAAGCGGGCAGTGAGCGCAACGCAATTAATGTAAGTTAGCTCACTCATTAGGCACCGGGATCTCGACCGATGCCCTTGAGAGCCTTCAACCCAGTGCCACCTGACGTCTAAGAAAAGGAATATTCAGCAATTTGCCCGTGCCGAAGAAAGGCCCACCCGTGAAGGTGAGCCAGTGAGTTGATTGCTACGTAAATAAATGTGAGCGGATAACAATTGACATTGTGAGCGGATAACAAGATACTGAGCACATCAGCATTGCATCTTAATCTAGCAGGGGAGTCTTTATGGAGCAAAACCCGCAGTCACAGCTGAAACTTCTTGTCACCCGTGGTAAGGAGCAAGGCTATCTGACCTATGCCGAGGTCAATGACCATCTGCCGGAAGATATCGTCGATTCAGATCAGATCGAAGACATCATCCAAATGATCAACGACATGGGCATTCAGGTGATGGAAGAAGCACCGGATGCCGATGATCTGATGCTGGCTGAAAACACCGCGGACGAAGATGCTGCCGAAGCCGCCGCGCAGGTGCTTTCCAGCGTGGAATCTGAAATCGGGCGCACGACTGACCCGGTACGCATGTACATGCGTGAAATGGGCACCGTTGAACTGTTGACCCGCGAAGGCGAAATTGACATCGCTAAGCGTATTGAAGACGGGATCAACCAGGTTCAATGCTCCGTTGCTGAATATCCGGAAGCGATCACCTATCTGCTGGAACAGTACGATCGTGTTGAAGCAGAAGAAGCGCGTCTGTCCGATCTGATCACCGGCTTTGTTGACCCGAACGCAGAAGAAGATCTGGCACCTACCGCCACTCACGTCGGTTCTGAGCTTTCCCAGGAAGATCTGGACGATGACGAAGATGAAGACGAAGAAGATGGCGATGACGACAGCGCCGATGATGACAACAGCATCGACCCGGAACTGGCTCGCGAAAAATTTGCGGAACTACGCGCTCAGTACGTTGTAACGCGTGACACCATCAAAGCGAAAGGTCGCAGTCACGCTACCGCTCAGGAAGAGATCCTGAAACTGTCTGAAGTATTCAAACAGTTCCGCCTGGTGCCGAAGCAGTTTGACTACCTGGTCAACAGCATGCGCGTCATGATGGACCGCGTTCGTACGCAAGAACGTCTGATCATGAAGCTCTGCGTTGAGCAGTGCAAAATGCCGAAGAAAAACTTCATTACCCTGTTTACCGGCAACGAAACCAGCGATACCTGGTTCAACGCGGCAATTGCGATGAACAAGCCGTGGTCGGAAAAACTGCACGATGTCTCTGAAGAAGTGCATCGCGCCCTGCAAAAACTGCAGCAGATTGAAGAAGAAACCGGCCTGACCATCGAGCAGGTTAAAGATATCAACCGTCGTATGTCCATCGGTGAAGCGAAAGCCCGCCGTGCGAAGAAAGAGATGGTTGAAGCGAACTTACGTCTGGTTATTTCTATCGCTAAGAAATACACCAACCGTGGCTTGCAGTTCCTTGACCTGATTCAGGAAGGCAACATCGGTCTGATGAAAGCGGTTGATAAATTCGAATACCGCCGTGGTTACAAGTTCTCCACCTACGCAACCTGGTGGATCCGTCAGGCGATCACCCGCTCTATCGCGGATCAGGCGCGCACCATCCGTATTCCGGTGCATATGATTGAAACCATCAACAAGCTCAACCGTATTTCTCGCCAGATGCTGCAAGAGATGGGCCGTGAACCGACGCCGGAAGAACTGGCTGAACGTATGCTGATGCCGGAAGACAAGATCCGCAAAGTGCTGAAGATCGCCAAAGAGCCAATCTCCATGGAAACGCCGATCGGTGATGATGAAGATTCGCATCTGGGGGATTTCATCGAGGATACCACCCTCGAGCTGCCGCTGGATTCTGCGACCACCGAAAGCCTGCGTGCGGCAACGCACGACGTGCTGGCTGGCCTGACCGCGCGTGAAGCAAAAGTTCTGCGTATGCGTTTCGGTATCGATATGAACACCGACTACACGCTGGAAGAAGTGGGTAAACAGTTCGACGTTACCCGCGAACGTATCCGTCAGATCGAAGCGAAGGCGCTGCGCAAACTGCGTCACCCGAGCCGTTCTGAAGTGCTGCGTAGCTTCCTGGACGATTAATAATAAAGGAATTGAAACAGCGTATCTGGGATTCAAAAATTAGCAGAAAGTCAAAAGCCTCCGACCGGAGGCTTTTGACTAAAACTTCCCTTGGGGTTATCATTGGGGCTCACTCAAAGGCGGTAATCAGATAAAAAAAATCCTTAGCTTTCGCTAAGGATGATTTCTGCTAGTATTATTATTTGCCGACTACCTTGGTGATCTCGCCTTTCACGTAGTGGACAAATTCTTCCAACTGATCTGCGCGCGAGGCCAAGCGATCTTCTTCTTGTCCAAGATAAGCCTGTCTAGCTTCAAGTATGACGGGCTGATACTGGGCCGGCAGGCGCTCCATTGCCCAGTCGGCAGCGACATCCTTCGGCGCGATTTTGCCGGTTACTGCGCTGTACCAAATGCGGGACAACGTAAGCACTACATTTCGCTCATCGCCAGCCCAGTCGGGCGGCGAGTTCCATAGCGTTAAGGTTTCATTTAGCGCCTCAAATAGATCCTGTTCAGGAACCGGATCAAAGAGTTCCTCCGCCGCTGGACCTACCAAGGCAACGCTATGTTCTCTTGCTTTTGTCAGCAAGATAGCCAGATCAATGTCGATCGTGGCTGGCTCGAAGATACCTGCAAGAATGTCATTGCGCTGCCATTCTCCAAATTGCAGTTCGCGCTTAGCTGGATAACGCCACGGAATGATGTCGTCGTGCACAACAATGGTGACTTCTACAGCGCGGAGAATCTCGCTCTCTCCAGGGGAAGCCGAAGTTTCCAAAAGGTCGTTGATCAAAGCTCGCCGCGTTGTTTCATCAAGCCTTACGGTCACCGTAACCAGCAAATCAATATCACTGTGTGGCTTCAGGCCGCCATCCACTGCGGAGCCGTACAAATGTACGGCCAGCAACGTCGGTTCGAGATGGCGCTCGATGACGCCAACTACCTCTGATAGTTGAGTCGATACTTCGGCGATCACCGCTTCCCTCATGATGTTTAACTTTGTTTTAGGGCGACTGCCCTGCTGCGTAACATCGTTGCTGCTCCATAACATCAAACATCGACCCACGGCGTAACGCGCTTGCTGCTTGGATGCCCGAGGCATAGACTGTACCCCAAAAAAACAGTCATAACAAGCCATGAAAACCGCCACTGCGCCGTTACCACCGCTGCGTTCGGTCAAGGTTCTGGACCAGTTGCGTGAGCGCATAACACCCCTTGTATTACTGTTTATGTAAGCAGACAGTTTTATTGTTCATGATGATATATTTTTATCTTGTGCAATGTAACATCAGAGATTTTGAGACACAACGTGGCTTTGTTGAATAAATCGAACTTTTGCTGAGTTGAAGGATCAGGTTACATTGTCGATCTGTTCATGGTGAACAGCTTTAAATGCACCAAAAACTCGTAAAAGCTCTGATGTATCTATCTTTTTTACACCGTTTTCATCTGTGCATATGGACAGTTTTCCCTTTGATATCTAACGGTGAACAGTTGTTCTACTTTTGTTTGTTAGTCTTGATGCTTCACTGATAGATACAAGAGCCATAAGAACCTCAGATCCTTCCGTATTTAGCCAGTATGTTCTCTAGTGTGGTTCGTTGTTTTTGCGTGAGCCATGAGAACGAACCATTGAGATCATACTTACTTTGCATGTCACTCAAAAATTTTGCCTCAAAACTGGTGAGCTGAATTTTTGCAGTTAAAGCATCGTGTAGTGTTTTTCTTAGTCCGTTATGTAGGTAGGAATCTGATGTAATGGTTGTTGGTATTTTGTCACCATTCATTTTTATCTGGTTGTTCTCAAGTTCGGTTACGAGATCCATTTGTCTATCTAGTTCAACTTGGAAAATCAACGTATCAGTCGGGCGGCCTCGCTTATCAACCACCAATTTCATATTGCTGTAAGTGTTTAAATCTTTACTTATTGGTTTCAAAACCCATTGGTTAAGCCTTTTAAACTCATGGTAGTTATTTTCAAGCATTAACATGAACTTAAATTCATCAAGGCTAATCTCTATATTTGCCTTGTGAGTTTTCTTTTGTGTTAGTTCTTTTAATAACCACTCATAAATCCTCATAGAGTATTTGTTTTCAAAAGACTTAACATGTTCCAGATTATATTTTATGAATTTTTTTAACTGGAAAAGATAAGGCAATATCTCTTCACTAAAAACTAATTCTAATTTTTCGCTTGAGAACTTGGCATAGTTTGTCCACTGGAAAATCTCAAAGCCTTTAACCAAAGGATTCCTGATTTCCACAGTTCTCGTCATCAGCTCTCTGGTTGCTTTAGCTAATACACCATAAGCATTTTCCCTACTGATGTTCATCATCTGAGCGTATTGGTTATAAGTGAACGATACCGTCCGTTCTTTCCTTGTAGGGTTTTCAATCGTGGGGTTGAGTAGTGCCACACAGCATAAAATTAGCTTGGTTTCATGCTCCGTTAAGTCATAGCGACTAATCGCTAGTTCATTTGCTTTGAAAACAACTAATTCAGACATACATCTC |
| Bujard RBS | Reporter Plasmid RBS for sfGFP | GAATTCATTAAAGAGGAGAAAGGT |
| sfGFP | Fluorescent Reporter for Reporter Plasmid | ATGCGTAAAGGCGAAGAGCTGTTCACTGGTTTCGTCACTATTCTGGTGGAACTGGATGGTGATGTCAACGGTCATAAGTTTTCCGTGCGTGGCGAGGGTGAAGGTGACGCAACTAATGGTAAACTGACGCTGAAGTTCATCTGTACTACTGGTAAACTGCCGGTACCTTGGCCGACTCTGGTAACGACGCTGACTTATGGTGTTCAGTGCTTTGCTCGTTATCCGGACCACATGAAGCAGCATGACTTCTTCAAGTCCGCCATGCCGGAAGGCTATGTGCAGGAACGCACGATTTCCTTTAAGGATGACGGCACGTACAAAACGCGTGCGGAAGTGAAATTTGAAGGCGATACCCTGGTAAACCGCATTGAGCTGAAAGGCATTGACTTTAAAGAAGACGGCAATATCCTGGGCCATAAGCTGGAATACAATTTTAACAGCCACAATGTTTACATCACCGCCGATAAACAAAAAAATGGCATTAAAGCGAATTTTAAAATTCGCCACAACGTGGAGGATGGCAGCGTGCAGCTGGCTGATCACTACCAGCAAAACACTCCAATCGGTGATGGTCCTGTTCTGCTGCCAGACAATCACTATCTGAGCACGCAAAGCGTTCTGTCTAAAGATCCGAACGAGAAACGCGATCACATGGTTCTGCTGGAGTTCGTAACCGCAGCGGGCATCACGCATGGTATGGATGAACTGTACAAATAA |
| apFAB71 | Original Promoter for Reporter Plasmid | TTGACATCGCATCTTTTTGTACCTATAATAGATTCATGATGA |
| apFAB71-∆(-7) | -35 Dependent Promoter for Reporter Plasmid | TTGACATCGCATCTTTTTGTACCTATAAAGATTCATGATGA |
| TTCATC-apFAB71-∆(-7) | Target Promoter | TTCATCATCGCATCTTTTTGTACCTATAAAGATTCATGATGA |
| GGAACC-apFAB71-∆(-7) | Target Promoter | GGAACCATCGCATCTTTTTGTACCTATAAAGATTCATGATGA |
| CCGCCG-apFAB71-∆(-7) | Target Promoter | CCGCCGATCGCATCTTTTTGTACCTATAAAGATTCATGATGA |
| CCCCTC-apFAB71-∆(-7) | Target Promoter | CCCCTCATCGCATCTTTTTGTACCTATAAAGATTCATGATGA |
| GCTACC-apFAB71-∆(-7) | Target Promoter | GCTACCATCGCATCTTTTTGTACCTATAAAGATTCATGATGA |
